# Mechanistic insight into substrate processing and allosteric inhibition of human p97

**DOI:** 10.1101/2021.02.08.430295

**Authors:** Man Pan, Yuanyuan Yu, Huasong Ai, Qingyun Zheng, Yuan Xie, Lei Liu, Minglei Zhao

**Affiliations:** Department of Biochemistry and Molecular Biology, The University of Chicago; Chicago, IL 60637, USA.; Tsinghua-Peking Center for Life Sciences, Department of Chemistry, Tsinghua University; Beijing 100084, China.

**Author notes:** Corresponding authors: Minglei Zhao.; Lei Liu.; Man Pan. These authors contributed equally to the work (Man Pan, Yuanyuan Yu).

**Keywords:** p97, NMS-873, single-particle cryo-EM, translocation, allosteric inhibition

## Abstract

p97 processes ubiquitinated substrates and plays a central role in cellular protein homeostasis. Previous studies have showed that it is a potential drug target for cancer, neurodegenerative disease, and viral infections. Here, we report a series of cryo-electron microscopy (cryo-EM) structures of substrate-engaged human p97 complex that captured “power stroke”-like motions of both the D1 and D2 ATPase rings of p97. A key feature of these structures is the critical conformational changes of the inter-subunit signaling (ISS) motifs, which tightens the binding of nucleotides and neighboring subunits, and contributes to the spiral staircase conformation of the D1 and D2 rings. We further determined the cryo-EM structure of human p97 in complex with NMS-873, the most potent p97 inhibitor. The structures showed that NMS-873 binds at a cryptic groove in the D2 domain and interacts with the ISS motif, preventing its conformational change, thus blocking substrate translocation allosterically.

## INTRODUCTION

p97 is a 540 kDa protein that belongs to the family of ATPases Associated with diverse cellular Activities (AAA)^1–3^. It undergoes repetitive ATP hydrolysis and conformational changes to extract ubiquitinated substrates from various cellular components, such as membranes, ribosomes, and chromatin^4^. p97 plays a central role in many cellular processes, including endoplasmic reticulum-associated degradation (ERAD), mitochondrial-associated degradation, chromatin-associated degradation, autophagy, and endosomal trafficking^5^. Mutations in human p97 have been found in multisystem proteinopathy (MSP), a dominantly inherited degenerative disorder that can affect muscle, bone and the central nervous system, that can manifest clinically as inclusion body myopathy with early-onset Paget disease and frontotemporal dementia (IBMPFD) and familial amyotrophic lateral sclerosis (ALS)^6, 7^. Moreover, recent studies have also targeted p97 for the treatment of cancer and viral infections^8, 9^.

p97 has two tandem AAA domains named D1 and D2 and an additional N domain at the N-terminus (**Fig. 1a**). This protein functions as a homomeric hexamer, with twelve copies of ATPase domains organized into two rings^10^. Crystal structures of full-length p97 were determined more than a decade ago^11, 12^. High-resolution single-particle cryo-electron microscopy (cryo-EM) structures have also been reported^13, 14^. However, all the structures of p97 determined thus far adopt a sixfold symmetric conformation, providing limited information regarding the mechanism since almost all AAA proteins, including close homologues of p97, break symmetry when engaged with substrates^15–17^. Recent cryo-EM studies of Cdc48, the yeast homolog of p97, in complex with either in vivo^18^ or in vitro^19^ substrates provided crucial insight into the working states. However, with medium resolution (3.7 Å to 4.2 Å) and limited conformational states, critical mechanistic questions, such as how the AAA domains in two rings coordinate to achieve translocation, remain to be answered. In addition, owing to its biomedical importance, many small-molecule inhibitors of p97 have been developed in the past decade^8, 20, 21^. NMS-873, a non-competitive allosteric inhibitor of p97^21^, has been shown to be active in tumor cell lines and IBMPFD patient fibroblasts^21, 22^. More importantly, low nanomolar concentrations of NMS-873 was recently reported to inhibit the replication of several viruses, including severe acute respiratory syndrome coronavirus 2 (SARS-CoV-2) and influenza virus, through its inhibition of p97^23–25^. Little is known regarding the allosteric inhibition mechanism due to the lack of a substrates-engaged human p97 structure in a working state, which is of considerable biological and clinical interest.

**Fig. 1:**
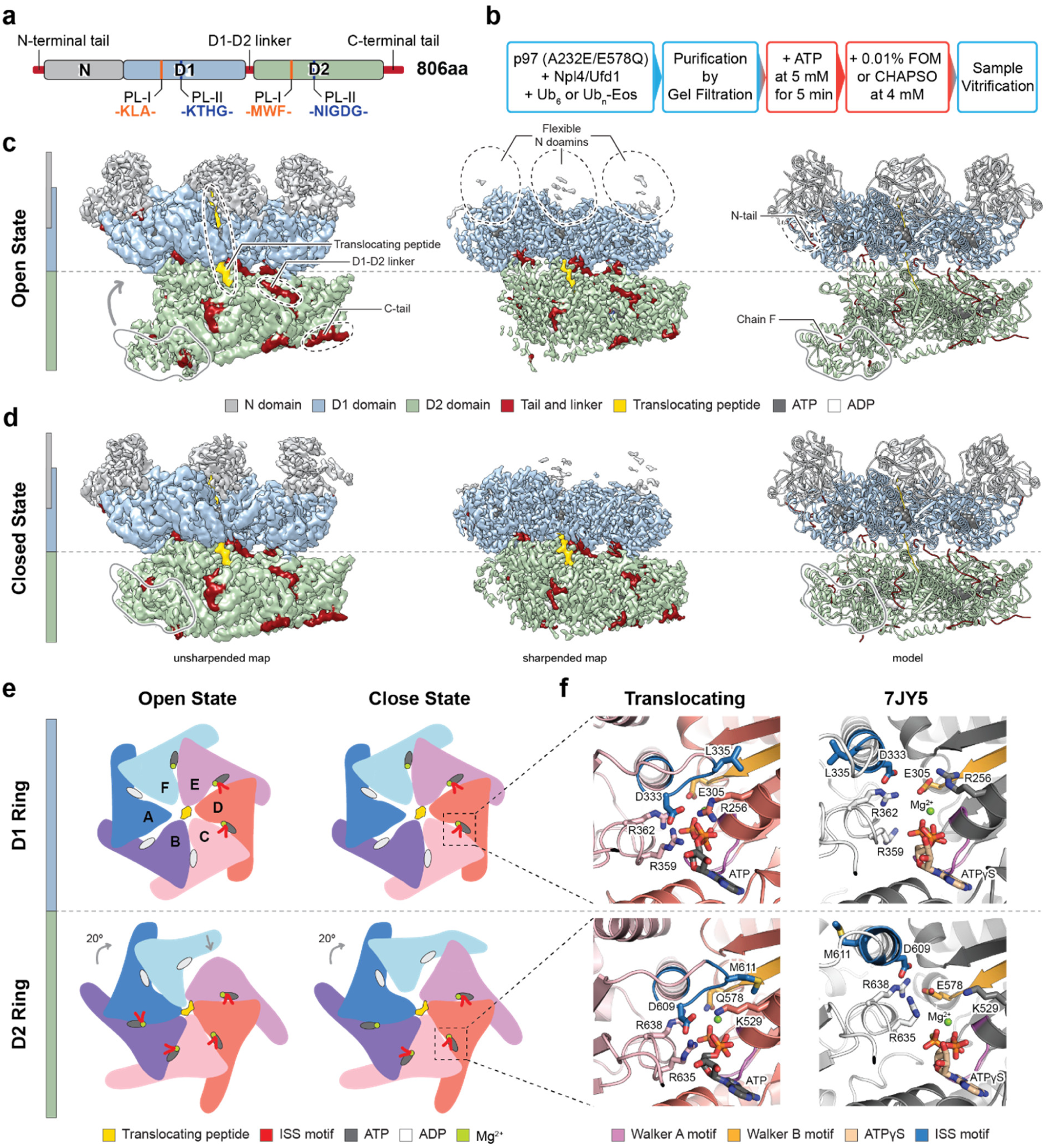
Structures of human p97 in working states. **a,** Domain architecture of human p97. The color code of individual domains is used whenever possible. Pore loop residues in D1 and D2 domains are listed. **b,** Sample preparation procedures. Steps performed at 4 °C are in blue boxes. Steps performed at room temperature are in red boxes. **c** and **d,** Cryo-EM maps and models of human p97 engaged with a translocating peptide. The maps and models are aligned. Contour level: unsharpened map, 0.015; sharpened map, 0.035. **c,** Open state. **d,** Closed state. **e,** Illustrations showing the arrangement of the D1 and D2 rings in the open and closed states, respectively. **f,** Magnified views of compressed ATP binding sites in the D1 and D2 rings. The corresponding ATPγS binding sites from the nontranslocating structure (PDB ID: 7JY5) are shown for comparison.

In this study, starting from a fully assembled human p97 complex with the cofactor Npl4/Ufd1 and a substrate, we triggered ATP hydrolysis and determined a series of substrate-engaged p97 structures with resolutions ranging from 2.9 to 3.8 Å using single-particle cryo-EM. The structures captured ATP hydrolysis driven “power stroke”-like motions of both the D1 and D2 rings of p97 and elucidated how the two rings coordinate to achieve translocation through inter-subunit signaling (ISS) motifs, the pore loops, and N- and C-terminal tails. We further determined the cryo-EM structure of p97 in the presence of NMS-873 at a resolution of 2.4 Å and uncovered a novel inhibition mechanism. Finally, using NMS-873 at a substoichiometric concentration, we captured a series of intermediate states, suggesting how p97 coordinates with its cofactor Npl4 to initiate substrate translocation. Taken together, our results established a comprehensive understanding of substrate processing and allosteric inhibition of human p97.

## RESULTS

### Capture the working states of human p97

First, we assembled complexes of p97 from recombinantly overexpressed components without introducing additional ATP. Two mutations, A232E and E578Q, were introduced into the human p97 construct to stabilize the complex as previously described ^26^. The A232E mutation has been found in MSP patients and has been reported as an activating mutant with its N domains shifted toward the “up conformation” ^27^. E578Q is a mutation in the Walker B motif of the D2 domain that slows substrate unfolding^19, 28^. Combined, the two mutations decreased but maintained the unfolding activity of p97 (**Extended Data Fig. 1a**) and minimized the conformational heterogeneity resulting from the N domains. All the structural work in this study used the p97 (A232E/E578Q) mutant and Npl4/Ufd1 as the cofactors. For substrates, either polyubiquitinated Ub-Eos (Ub_n_-Eos) or K48-linked hexa-ubiquitin (Ub_6_) was used. The assembled complex, named p97-Npl4/Ufd1-Ub_n_-Eos or p97-Npl4/Ufd1-Ub_6_, eluted as single peaks in the size-exclusion chromatograms (**Extended Data Fig. 1b and c**). To capture the working states of p97, ATP hydrolysis was triggered by adding 5 mM ATP to the purified complexes followed by incubation at room temperature for 5 minutes. To solve the orientation preference problem to the greatest extent, either CHAPSO or fluorinated octyl maltoside (FOM) was added immediately before sample vitrification (**Fig. 1b**). Three datasets, namely, p97-Npl4/Ufd1-Ub_n_-Eos-CHAPSO, p97-Npl4/Ufd1-Ub_n_-Eos-FOM, and p97-Npl4/Ufd1-Ub_6_-FOM, were collected, and a total of 9 different cryo-EM maps with resolutions ranging from 2.9 to 3.8 Å were reconstructed using single-particle analysis (**Extended Data Fig. 2-4, Supplementary Table 1**). Atomic models were built based on existing p97 structures (**Supplementary Table 2**). For well-resolved regions, individual residues were manually built into the density. For flexible regions, rigid body docking was used to fit individual secondary structure elements or subdomains (see Methods for details).

### Open and closed states of the D2 ring

In the p97-Npl4/Ufd1-Ub_n_-Eos-CHAPSO dataset, two conformational states of the D2 ring were resolved (**Fig. 1c** and **d**). In the open state (3.6 Å, **Fig. 1c**, corresponding to class 2 in **Extended Data Fig. 2**), all six D2 domains interacted with the substrate and formed an open, right-handed spiral staircase, whereas in the closed state (3.1 Å, **Fig. 1d**, corresponding to class 1 in **Extended Data Fig. 2**), the bottom D2 domain (termed chain F) detached from the substrate and shifted up to interact with the top D2 domain (termed chain E), closing the gap of the spiral in the open state. Only five D2 domains (chains A-E) interacted with the substrate in the closed state. The two states suggested a “power stroke”-like motion of the D2 domain (**Supplementary Movie 1**). Compared to the same complex structure in the presence of ATPγS that was previously published^26^ (PDB ID: 7JY5), there was a relative 20 degree rotation of the D2 ring to the D1 ring in both states (**Fig. 1e** and **Extended Data Fig. 6a**). A superimposition of the two states showed that the D2 domain immediately above chain F (chain A) also shifted up from the open state to the closed state (**Extended Data Fig. 7a** and **Supplementary Movie 1**). The particles in the open state were significantly less than those in the closed state (13% vs. 70%), suggesting that the open state is relatively transient. A third class was resolved from the 3D classification (3.8 Å, class 3 in **Extended Data Fig. 2**), with better resolved density for the cofactor Npl4 but otherwise the same conformation as the closed state.

### Conformational changes in the D1 ring

In the p97-Npl4/Ufd1-Ub_n_-Eos-FOM dataset, three conformational states of the D1 ring were resolved (**Extended Data Fig. 3**). All the D2 rings were in the same conformation as the closed state described above. Using the same naming convention, we discovered from the superimposed structures that chain A shifted up slightly from class 1 (3.6 Å) to class 2 (3.4 Å) and much further from class 2 to class 3 (3.0 Å) (**Extended Data Fig. 8a**). No open spiral conformation similar to the open state of the D2 ring was observed for the D1 ring, although a right-handed spiral staircase was still formed by chains B-F. The three states of the D1 ring also suggested a “power stroke”-like motion despite the differences from the D2 ring (**Supplementary Movie 2**). It is worth noting that no conformational changes in the D1 ring were observed for the p97-Npl4/Ufd1-Ub_n_-Eos-CHAPSO dataset. The conformation of the D1 ring in the open and closed states was very similar to class 3 of the p97-Npl4/Ufd1-Ub_n_-Eos-FOM dataset (see Discussion).

### Sequential but asynchronous hydrolysis in D1 and D2 rings

The resolutions of the maps allowed us to build atomic models and identify the nucleotides in most nucleotide binding sites (**Extended Data Fig. 7d and S8d**). For the less well resolved chains A and F, the identity of the nucleotides could not be determined with certainty, but the density of a nucleotide was clearly visible (**Extended Data Fig. 7d and S8d**). All nucleotide binding sites were occupied, suggesting that the nucleotide exchange might be very fast. When superimposing all the AAA domains within the same ring, two major conformations were observed: one with a larger angle between the α/β and the α subdomain, potentially corresponding to an ATP binding conformation, and the other with a smaller angle between the α/β and the α subdomain, potentially corresponding to an ADP binding conformation (**Extended Data Fig. 7c and 8c**). Such binary conformations have been observed for other substrate-engaged AAA+ structures ^15–19, 29, 30^.

When focusing on the well-resolved sites, we discovered a new compact ATP binding mode in both the D1 and D2 rings involving a previously defined ISS motif^31^. Specifically, the ISS motif changed from a helical conformation to a triangular loop that inserted into the neighboring nucleotide binding site (**Fig. 1e** and **f**), and these conformational changes were closely related to the spiral staircases formed by the pore loops (**Fig. 2a** and **Extended Data Fig. 5a and b**). L335 in the D1 domain and M611 in the D2 domain are the key residues interacting with the hydrophobic residues in the Walker B motifs of the neighboring subunits (**Fig. 1f**). As a result, ATP is coordinated by three more residues than the ATPγS-bound structure (PDB ID: 7JY5), including one additional arginine finger residue (R362 in D1 and R638 in D2), an aspartic acid from the ISS motif (D333 in D1 and D609 in D2), and a basic residue following the Walker A motif (R256 in D1 and K529 in D2) (**Fig. 1f**). This binding mode was observed only when ATP was engaged and only for two to four consecutive binding sites in both rings (**Fig. 1e**, **Extended Data Fig. 7b and S8b**). We hypothesize that this tight binding mode is required for the formation of spiral staircases and the rotational compression of ATPase rings, both essential for substrate translocation. For ATP hydrolysis or ADP release to occur, the ISS motif needs to be retracted to loosen the binding with the nucleotide. For example, the ISS motif in the D2 domain of chain A retracted in the closed state (not in the open state) since the neighboring D2 domain in chain B is the next subunit to undergo ATP hydrolysis or ADP release (**Fig. 1e** and **Extended Data Fig. 7b**).

**Fig. 2:**
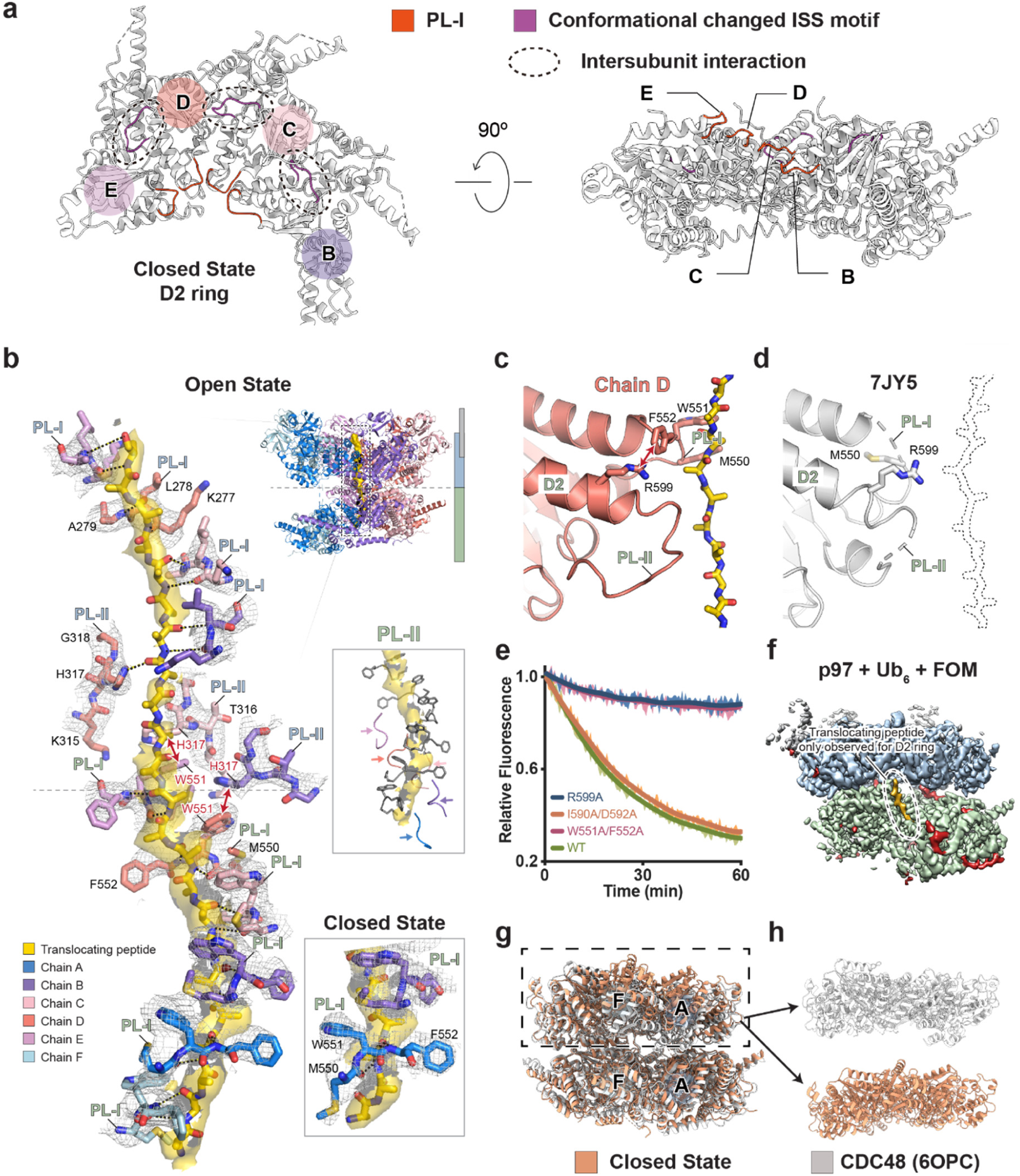
Interactions between the pore loops and the substrate. **a,** Illustrations showing the arrangement of chains B-E in the D2 ring of the closed state. **b,** Interactions between the translocating peptide and the interacting pore loops in the D1 and D2 rings. The density of the pore loops and the translocating peptide are shown as a gray mesh and yellow surface, respectively. Contour level: 6.0 root-mean-square deviation (rmsd). The closed state and PL-II of the D2 ring are shown in the insets. Hydrogen bonds between the pore loops and the peptide are labeled with black dotted lines. π-π interactions between H317 and W551 are labeled with red double arrows. **c and d,** A comparison of the D2 domain in chain D of the open state (panel **c**) and that of the nontranslocating structure (panel **d**, PDB ID: 7JY5). The D2 domains are aligned. **e,** Substrate unfolding assay for the pore loop mutants of p97. The error bands represent the standard deviation from triplicate experiments. **f,** Unsharpened cryo-EM map (class 1) of the p97-Npl4/Ufd1-Ub_6_-FOM dataset (**Extended Data Fig. 4**). The map is shown in a similar orientation to Fig. 1d and 1e. Contour level: 0.015. **g**, A comparison of cryo-EM models of translocating structures of p97 (closed state, orange) and Cdc48 (PDB ID: 6OPC, light grey). The two models were aligned based on the D2 ring. **h**, A close view of the D1 rings in panel **g**.

Another unexpected observation is that the hydrolysis in the D1 and D2 rings is not in sync, which means that at a given time, the D1 and D2 domains of the same chain may engage different nucleotides. If viewed from the top, hydrolysis propagates counterclockwise in both ATPase rings, with the D1 ring one or two subunits ahead of the D2 ring (**Fig. 1e**). This feature is unique for human p97 and has not been observed for other type II AAA proteins, including yeast Cdc48.

### Interactions between the pore loops and the substrate

In all six maps discussed above, we observed the density corresponding to a translocating peptide going through the central pore of p97 (**Fig. 2b**). A continuous extended peptide could be modeled into the density. The peptide went through the pore from the N-terminus (bottom in **Fig. 2b**) to the C-terminus (top in **Fig. 2b**), as the other direction fit much worse into the density. This directionality of translocating peptide is consistent with that of Cdc48^18, 19^. The density corresponding to the side chains was averaged, and polyalanine was modeled.

In general, the density of the translocating peptide in the D2 ring was better than that in the D1 ring. Three residues in pore loop-I (PL-I) of the D2 domain, namely, M550, W551, and F552, interacted with the peptide like a “pitching grip” (**Fig. 2b**). Each PL-I formed two hydrogen bonds with the nitrogen and carbonyl oxygen of one residue in the translocating peptide through the carbonyl oxygen of M550 and nitrogen of F552. Every other residue in the translocating peptide was hydrogen bonded by a PL-I so 12 residues were engaged with six PL-I in the D2 ring, which formed a spiral in the open state, with chain F at the bottom and chain E at the top (**Fig. 2b**). In the closed state, the PL-I of chain F detached from the translocating peptide. The remaining five PL-I formed the spiral in a very similar way except for the side chain of M550 in chain A (now the bottom chain), which was pointing downwards and would not allow an additional PL-I to engage (**Fig. 2b** **inset**). In both the open and closed states, the density of five pore loop-II (PL-II) in the D2 ring was resolved (chains A to E) but at lower resolution (**Fig. 2b** **inset**). PL-II in the D2 ring did not form hydrogen bonds with the translocating peptide and likely played a less important role in translocation. We mutated the residues of PL-I and PL-II and performed the substrate unfolding assay^1, 26, 32^. Indeed, the PL-I mutation (W551A/F552A) abolished the unfolding activity, whereas the PL-II mutation (I590A/D592A) showed no effect (**Fig. 2e**). A closer examination revealed that R599 was π-stacked with F552 of PL-I (**Fig. 2c**). This interaction might be critical to stabilize the “pitching grip” conformation of PL-I since the R599A mutant did not show any unfolding activity, similar to the PL-I mutation (**Fig. 2e**). In contrast, the pore loops in the D2 domain of the nontranslocating structure (PDB ID: 7JY5) were mostly unstructured (**Fig. 2d**).

PL-I in the D1 ring interreacted with the translocating peptide in the same fashion as that in the D2 ring, except that only four PL-I (chains B-E) were engaged and the three residues were K277, L278, and A279 (**Fig. 2b**). The “pitching grip” and hydrogen bonding pattern still existed, but the interaction was not as strong as that of the D2 ring since the average length of the hydrogen bonds was 3.9 Å in the D1 ring and 3.4 Å in the D2 ring. Two PL-II of the D1 ring (chains C and D) were involved in the interaction with the peptide by forming a hydrogen bond through H317 (**Fig. 2b**). Interestingly, H317 of chains B and C also stacked with W551 (PL-I of D2) of chains D and E, respectively (**Fig. 2b**, red double arrow). This π-π stacking might be involved in the coordination of two ATPase rings.

To further investigate the behavior of p97 with substrates of different sizes, we used a much smaller substrate, Ub_6_ (**Extended Data Fig. 1c**), and collected a third dataset, p97-Npl4/Ufd1-Ub_6_-FOM (**Extended Data Fig. 4**). Again, three conformations of the D1 ring that were very similar to those of the p97-Npl4/Ufd1-Ub_n_-Eos-FOM dataset were resolved. Intriguingly, the density corresponding to the translocating peptide was resolved only in the D2 ring (**Fig. 2f**). The D1 ring was essentially void of the translocating peptide despite adopting the same conformation. One possible explanation is that the D1 ring is engaged only when a “complicated” substrate such as Ub_n_-Eos is encountered.

We noticed that the D2 ring of the closed state is similar to those of Cdc48-ADP⋅BeFx structures ^18, 19^, all of which have five PL-I engaging with the translocating peptide. When the D2 rings from these structures were superimposed, it was obvious that the two rings of p97 are out of register, that is, the lowest D2 and D1 domains are not in the same chain (**Fig. 2g** and **Extended Data Fig. 5c**), whereas in Cdc48, the two rings essentially formed a single “split washer” (**Fig. 2h** and **Extended Data Fig. 5d**). Accordingly, the entire chain F (named based on Fig. 1) is flexible in previous Cdc48-ADP⋅BeFx structures, whereas in our structures the D2 domain of chain F and the D1 domain of chain A are the most flexible. These features are consistent with the asynchronous hydrolysis occurring in the D1 and D2 rings.

### N- and C-terminal tails and the linker between D1 and D2 rings

In addition to the ATPase rings, some of the N- and C-terminal tails (**Fig. 1a**) of p97 were also resolved in the cryo-EM maps. Both the resolved N-terminal (L12 to R22) and C-terminal (G767-S775) tails interacted with the neighboring subunit in a counterclockwise manner if viewed from the top (N-terminal tail, **Fig. 3a**) or a clockwise manner if viewed from the bottom (C-terminal tail, **Fig. 3c**). The N-terminal tail interacted with an acidic patch in the α subunit of the D1 domain (**Fig. 3b**). The C-terminal tail interacted with a groove in the α subunit of the D2 domain. Such interactions have not been described before and may play important roles in the coordination between different chains. Truncations of either the N- or C-terminal tails greatly affected the unfolding activity of p97 (**Fig. 3f**). The linker between the D1 and D2 rings was resolved for chains B-E. Compared to the structure of nontranslocating p97 (PDB ID: 7JY5), a 19 Å shift of L464 was observed (**Fig. 3e**). In the translocating structure, L464 interacted with A569 in the D2 domain, which contributed to the relative twist of the D2 ring (**Fig. 1e**). The L464A mutation decreased the unfolding activity of p97 (**Fig. 3f**). Notably, the N- and C-terminal tails and the linker regions are less conserved between human p97 and yeast Cdc48, which might contribute to the different conformations of the ATPase rings.

**Fig. 3:**
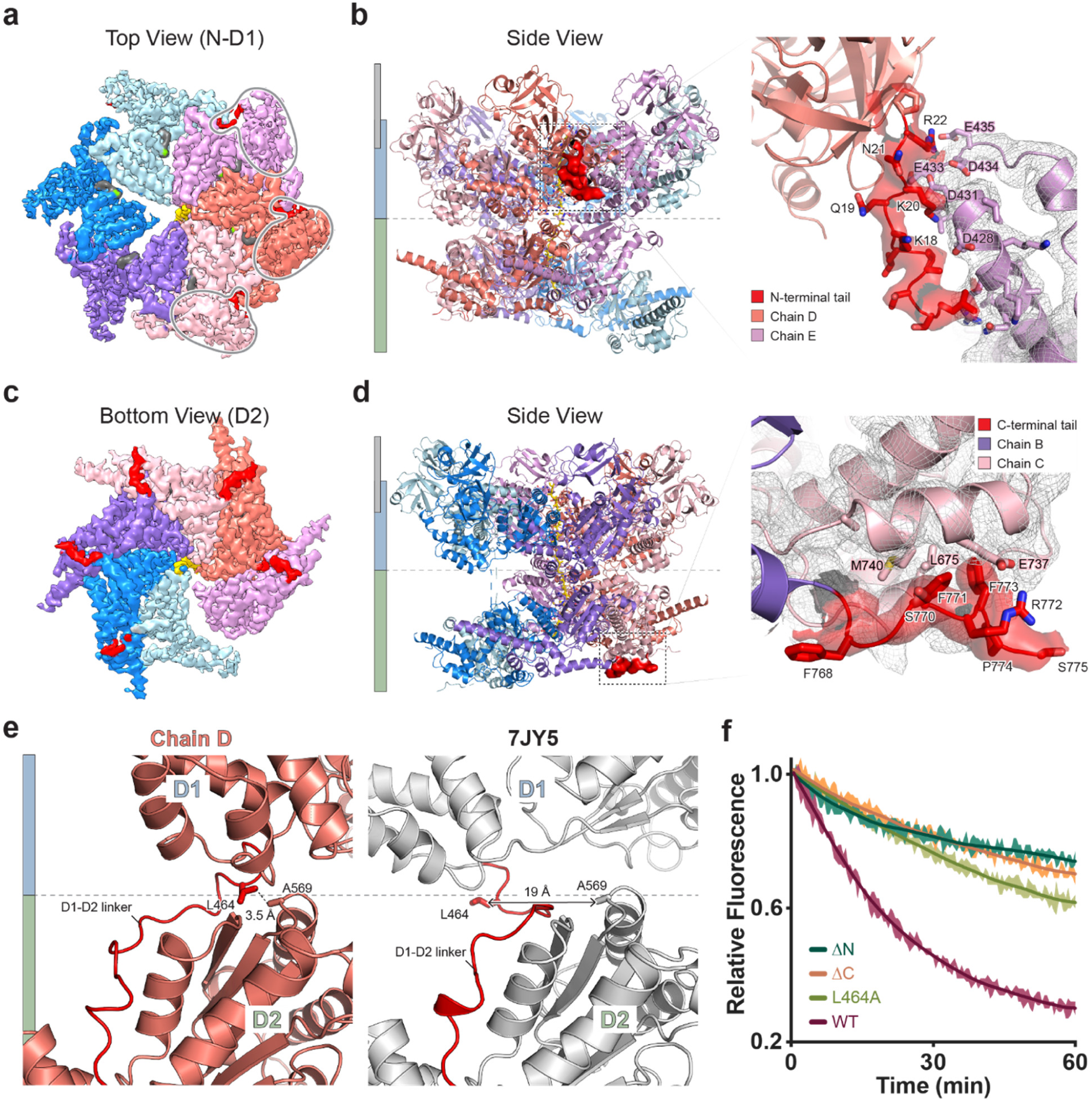
N- and C-terminal tails and the linker between the D1 and D2 rings. **a,** Top view of the unsharpened cryo-EM map focused on the N and D1 domains (open state). N-terminal tails are colored in red. Contour level: 0.015. **b,** A side view of the cryo-EM model of the open state with the density of one N-terminal tail highlighted and magnified. Contour level: 6.0 rmsd. **c,** A bottom view of the unsharpened cryo-EM map focused on the D2 ring (open state). C-terminal tails are shown in red. Contour level: 0.015. **d,** A side view of the cryo-EM model of the open state with the density of one C-terminal tail highlighted and magnified. Contour level: 6.0 rmsd. **e,** A comparison of the D1-D2 linker in chain D of the open state and that of the nontranslocating structure (PDB ID: 7JY5). The D2 domains are aligned. **f,** Substrate unfolding assay for tail and linker mutants of p97. The error bands represent the standard deviation from triplicate experiments.

### NMS-873 inhibits the translocation by locking the ISS motif

Inhibition of p97 has emerged as a promising strategy in anticancer and antiviral studies^8^. NMS-873, the first discovered allosteric inhibitor of p97 (**Fig. 4a**), was recently characterized as a candidate for the treatment of cancers, neurodegenerative diseases, and viral infections^22–24^. However, the mechanism of inhibition remained unclear. We introduced NMS-873 at saturation (80 μM) after triggering ATP hydrolysis and substrate translocation (**Fig. 4b**). Two datasets were collected using Ub_n_-Eos and Ub_6_ as the substrate. A single map was obtained from each dataset with overall resolutions of 2.4 Å (Ub_6_, **Extended Data Fig. 9**) and 2.8 Å (Ub_n_-Eos, **Extended Data Fig. 10**). The two maps are essentially identical; therefore, only the map with the higher resolution is shown (**Fig. 4c**). Unexpectedly, both maps showed that p97 maintained 6-fold symmetry and no translocating peptides were found in the central pore, although a substrate was present and ATP hydrolysis was triggered before adding NMS-873. In the 6-fold symmetric structure of p97, NMS-873 binds at a cryptic groove next to the ISS motif of the D2 domain (**Fig. 4d**) and is surrounded by many hydrophobic residues (**Fig. 4e**). A critical interaction was observed between K615 in the ISS motif and the biphenyl group of NMS-873 (**Fig. 4d**). As a result, the ISS motif is locked and prevented from essential conformational changes discovered in the translocating structures (**Fig. 4f**). The pore loops in the D2 domain of the NMS-873-bound structure were resolved but retracted from the pore compared to the translocating structure (**Extended Data Fig. 6e**). Together, these structural features make it impossible to form a spiral staircase by the PL-1 of D2 ring, which might explain why no translocating peptide was found in the pore.

**Fig. 4:**
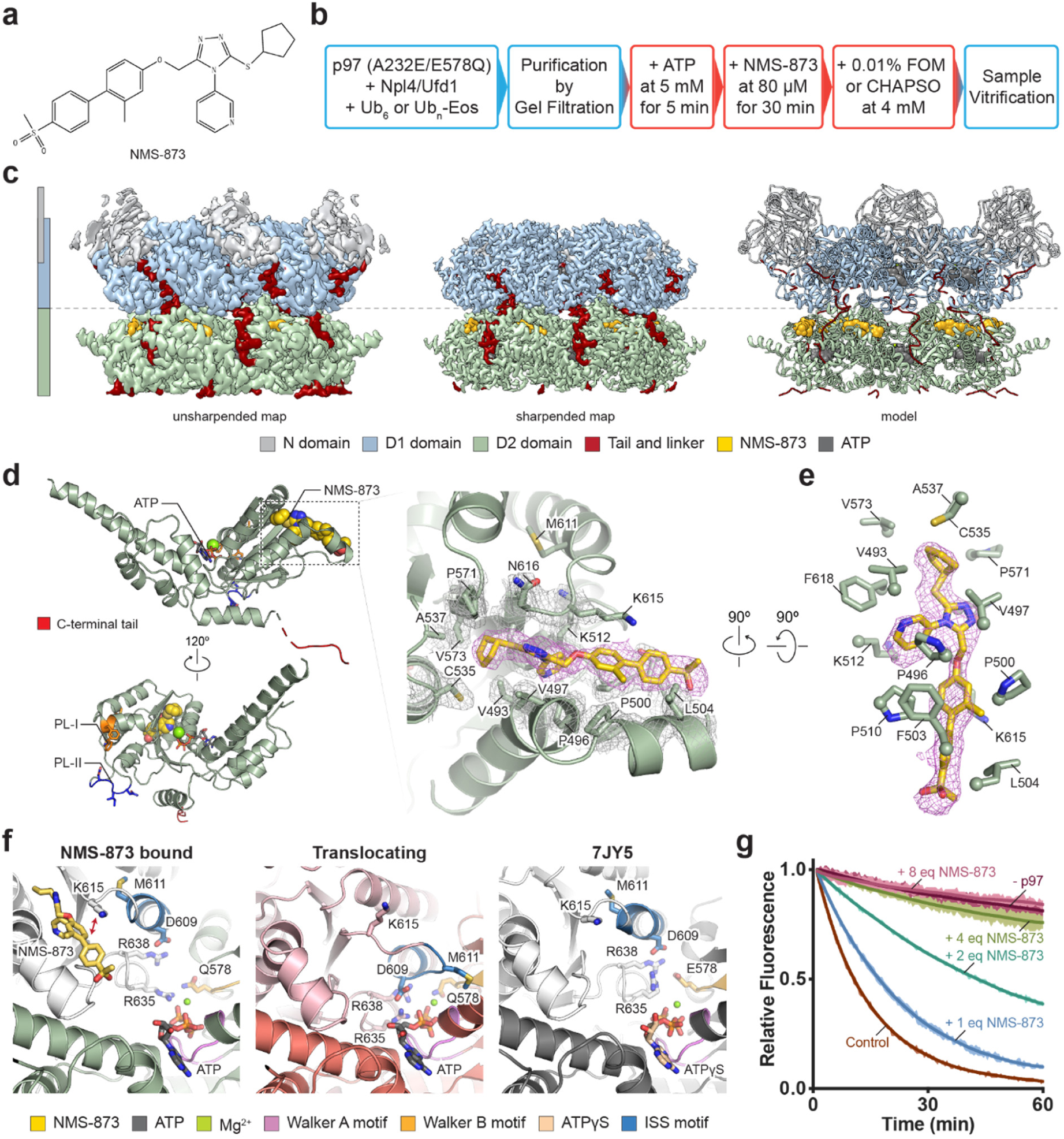
NMS-873 inhibits translocation by locking the ISS motif. **a,** Chemical structure of NMS-873. **b,** Sample preparation procedures. Steps performed at 4 °C are in blue boxes. Steps performed at room temperature are in red boxes. **c,** Cryo-EM maps and models of human p97 in complex with NMS-873. The maps and models are aligned. Contour level: unsharpened map, 0.018; sharpened map, 0.052. **d,** D2 domain of the complex structure in panel **c**, with the NMS-873 binding site zoomed in. The density of NMS-873 and the interacting residues are shown as magenta and gray mesh, respectively. Contour level: 6.0 rmsd. **e,** A different view of the NMS-873 binding site, with side chains of interacting residues shown. Contour level: 6.0 rmsd. **f,** A comparison of nucleotide binding sites in NMS-873 bound, translocating (chain D in open state), and nontranslocating (PDB ID: 7JY5) structures. The bottom D2 domains (green, salmon, and gray) are aligned. **g,** Substrate unfolding assay for wild-type p97 in the presence of different concentrations of NMS-873. One p97 hexamer can bind 6 equivalents (eq) of NMS-873.

Compared to the ATPγS-bound nontranslocating structure (PDB ID: 7JY5), the relative positions of the ATPase rings in the NMS-873-bound structure are similar (**Extended Data Fig. 6b**). The difference is that the bottom opening of the D2 ring is enlarged, which is similar to that of the ADP-bound structures^13^, although ATP was engaged (**Extended Data Fig. 6c**). The smaller angle between the α/β and the α subdomains of the D2 domains suggested that it is more similar to an ADP-bound state, indicating that NMS-873 can affect the conformation of the D2 ring regardless of the nucleotide states (**Extended Data Fig. 6d**). Superimposing the NMS-873-bound structure with another inhibitor-bound structure (UPCDC30245, PDB ID: 5FTJ ^13^) showed the similarity in the conformations of pore loops. The difference is that UPCDC30245 only shares part of the binding pocket of NMS-873 in the D2 domain (**Extended Data Fig. 6f and S6g**). Especially, the interaction between K615 and the biphenyl group of NMS-873 does not occur in the UPCDC30245-bound structure (**Fig. 4f** and **Extended Data Fig. 6g**), which was also reported from previous biochemical characterizations ^13, 21^. Combined with previous finding that UPCDC30245 binds p97 only in the presence of ADP and acts as an uncompetitive inhibitor, whereas NMS-873 can bind p97 in the absence of nucleotides and acts as a noncompetitive inhibitor ^8^, we concluded that the ISS-locking mechanism of NMS-873 through the interaction with K615 is the key to the non-competitive allosteric inhibition.

### Conformational states of Npl4 revealed by NMS-873 at a substoichiometric concentration

We further discovered that NMS-873 did not fully inhibit the unfolding activity of p97 at a substoichiometric concentration (6 equivalents (eq) is the stoichiometric concentration, **Fig. 4g**). On the reasoning that intermediate structures could be captured using NMS-873 as a modulator, a dataset was collected following the same protocol using Ub_n_-Eos as the substrate and in the presence of 10 μM NMS-873 (∼0.2 eq, **Fig. 5a**). As expected, multiple intermediate structures were resolved after 3D classification (**Extended Data Fig. 11**). Two out of six maps (∼11% particles) are in the nontranslocating state, with flat and symmetric ATPase rings and no density for a translocating peptide (**Fig. 6c**). One of the nontranslocating maps (Class 1, ∼8% particles) showed the density of NMS-873 (**Fig. 6c**, top) in all six binding pockets, suggesting that the binding of NMS-873 is highly cooperative. The cofactor Npl4 was resolved in four out of six maps (**Extended Data Fig. 11**). Both nontranslocating maps showed density for Npl4, with either one or two of its zinc finger motifs (ZF1 and ZF2, **Fig. 5b**) binding on top of the D1 ring (**Fig. 6c**). Such seesaw conformations of Npl4 have been described in a previous study^26^. The four translocating structures all had spiral ATPase rings and a translocating peptide similar to the conformation of the closed state (**Extended Data Fig. 11**). Two maps (class 5 and class 6) had very poor densities of Npl4 and differed in the D1 ring in a similar way to class 1 and class 3 of the p97-Npl4/Ufd1-Ub_n_-Eos-FOM dataset. The other two maps showed density for Npl4, but with its zinc finger motifs interacting with different chains (**Fig. 6d**), suggesting a rotation of Npl4 on top of the D1 ring. Although the resolution of Npl4 is not sufficient for atomic model building, a trace density connected to the translocating peptide can be seen to be engaged in a groove of Npl4 (**Fig. 6d**, top), which is reminiscent of a substrate processing complex structure of Cdc48^19^.

**Fig. 5:**
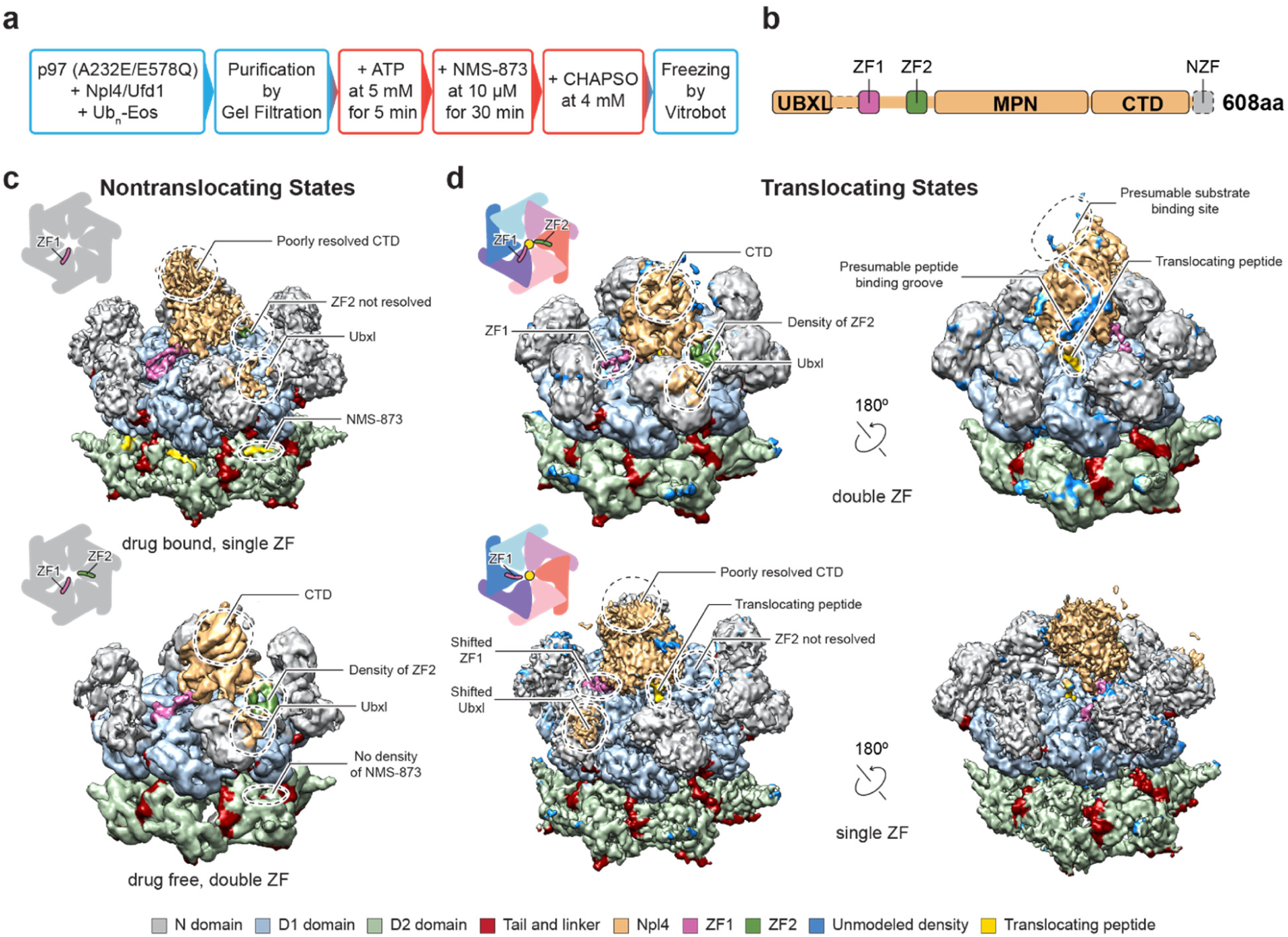
Conformational states of Npl4 in the presence of NMS873 at a substoichiometric concentration. **a,** Sample preparation procedures. Steps performed at 4 °C are in blue boxes. Steps performed at room temperature are in red boxes. **b,** Domain architecture of human Npl4. The color code of the individual motifs are the same as that in panels **c** and **d**. Dotted parts are not resolved in the cryo-EM maps. **c,** Two unsharpened cryo-EM maps in nontranslocating states resolved from the dataset (**Extended Data Fig. 10**), one bound with NMS-873 (top). Contour level: 0.013. **d,** Two unsharpened cryo-EM maps in translocating states resolved from the dataset. Contour level: top, 0.01; bottom, 0.013. Different positions of zinc finger motifs of Npl4 are illustrated in the insets.

## DISCUSSION

Here, we report the cryo-EM structures of human p97 in several working states, with a translocating peptide going through the central pore. Together, the structures revealed “power stroke”-like conformational changes in both the D1 and D2 rings. Surprisingly, the two AAA domains undergoing power stroke motion in each ring were not from the same chain, which seemed to be a unique feature of human p97 and was not observed for Cdc48 and other Type-II AAA proteins^15, 16, 18, 19, 29, 30^. The unusual structural feature corresponds to the asynchronous hydrolysis taking place in the two ATPase rings (**Fig. 1e** and **Extended Data Fig. 5c and d**). Regarding translocation, the D2 ring seemed to play a more important role by tightly interacting with the peptide through backbone hydrogen bonds. Previous biochemical studies have shown that the D2 ring of p97 had higher ATPase activity and provided the driving force for translocation, which is consistent with our structures^33^. The structures of Cdc48 also showed fewer interactions between the translocating peptide and the pore loops of the D1 ring^19^, further supporting this hypothesis. Regarding the role of the D1 ring in substrate translocation, we obtained some insights from the datasets using Ub_n_-Eos and Ub_6_ as the substrate, respectively. In the translocating structures with Ub_n_-Eos, peptide density was observed in both D1 and D2 rings, while in the structures with Ub_6_, the peptide density was only observed in the D2 ring. Therefore, D1 ring may be engaged when larger substrate proteins are encountered.

Although the structures strongly support a sequential hydrolysis model^34^, it is not clear whether the power strokes in the D1 and D2 rings take place concurrently. Multiple structures from a single dataset are needed to address this question. In this study, the motions of the D1 and D2 rings were resolved from two datasets, one using CHAPSO and the other using FOM as the detergent to fix the orientation preference. CHAPSO required a much higher concentration of the complex than FOM (8x) to keep the samples in the hole but seemed to preserve the complex better and yielded more particles with Npl4. Since the open state is a minor conformation (7% of the total particles, **Extended Data Fig. 2**), the higher concentration of the CHAPSO dataset may have contributed to its discovery.

One critical finding in this study is the conformational changes of the ISS motifs in both ATPase rings in the translocating structures (**Figs. 1F, 2a** and **Extended Data Fig. 5a-b**). Structures of human p97 in the presence of NMS-873 revealed the allosteric inhibition mechanism that NMS-873 binds and locks the ISS motifs in the D2 ring to prevent conformational changes (**Figs. 4d-f**), which further affects the pore loops and prevents substrate translocation. Targeting the ISS motif of p97, therefore, may serve as a potential strategy for the development of new inhibitors.

Multiple structures in the presence of NMS-873 at a substoichiometric concentration suggested that Npl4 may play a critical role in the initiation of translocation by unfolding the ubiquitin chain and threading it into the central pore through a seesaw motion. The pore loops in the D1 and D2 rings catch the thread-in peptide, and the two rings are compressed to spiral conformations. Following sequential ATP hydrolysis and power stroke motions circulating in both rings, the peptide in the pore is translocated downward. Npl4 might be unnecessary once the translocation starts.

Mutations in p97 have been linked to MSP^6, 35^. Structures of p97 in working states provided crucial insights into the consequences of the mutations (**Supplementary Table 3, Supplementary Movie 3**). The resolutions of the cryo-EM maps in this study are among the highest of type II AAA proteins^15, 17, 19, 30, 36^, allowing a comprehensive understanding of substrate processing and allosteric inhibition of human p97 and shedding light on the mechanism of other type II AAA proteins.

## ACKNOWLEDGEMENTS

We thank staff at the National Cryo-Electron Microscopy Facility at the Frederick National Laboratory and the Advanced Electron Microscopy Facility at the University of Chicago for the help in cryo-EM data collection.

## AUTHOR CONTRIBUTIONS

M.P., M.Z. and L.L. designed all the experiments and interpreted the results. M.P., Y.Y., H.A., Q.Z., and Y.X. cloned, expressed, and purified all the complexes and carried out related biochemical characterizations. M.P., Y.Y., and M.Z. performed cryo-EM data collection and processing. M.Z., M.P., and L.L. wrote the paper. M.Z., L.L., and M.P. supervised the project.

## EXTENDED DATA Figures and Legends

**Extended Data Fig. 1:**
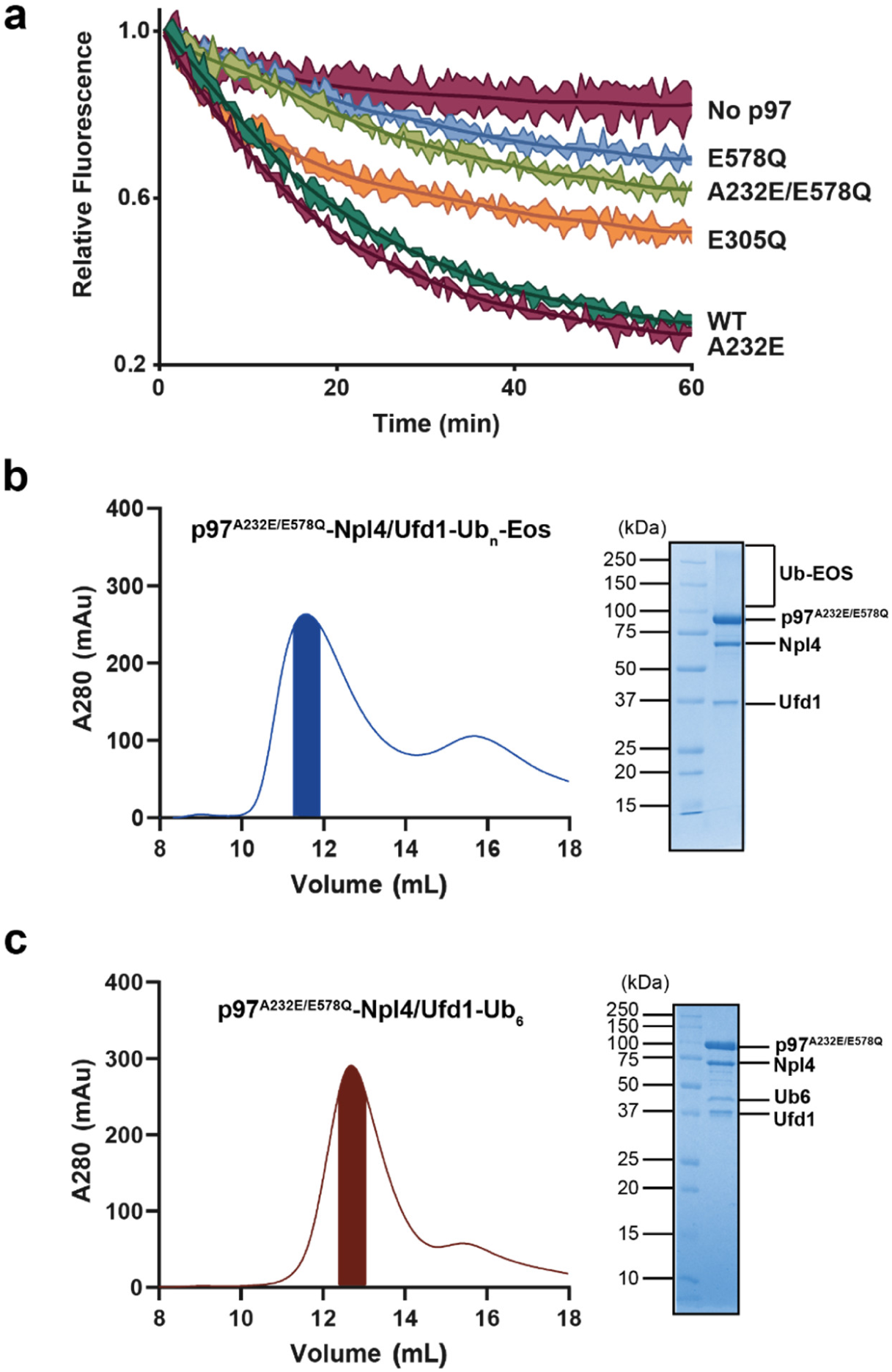
Substrate unfolding activity of various p97 mutants and the purification of cofactor- and substrate-engaged p97 complexes. **a,** Substrate unfolding assay for wild-type (WT) p97 and various mutations. E305Q and E578Q are the two Walker B mutations in the D1 and D2 domains, respectively. A232E is a disease mutation found in multisystem proteinopathy (MSP)^27, 32^. **b,** Gel filtration chromatogram and SDS-PAGE gel of the p97^A232E/E578Q^-Npl4/Ufd1-Ub_n_-Eos complex. **c,** Gel filtration chromatogram and SDS-PAGE gel of the p97^A232E/E578Q^-Npl4/Ufd1-Ub_n_-Eos complex.

**Extended Data Fig. 2:**
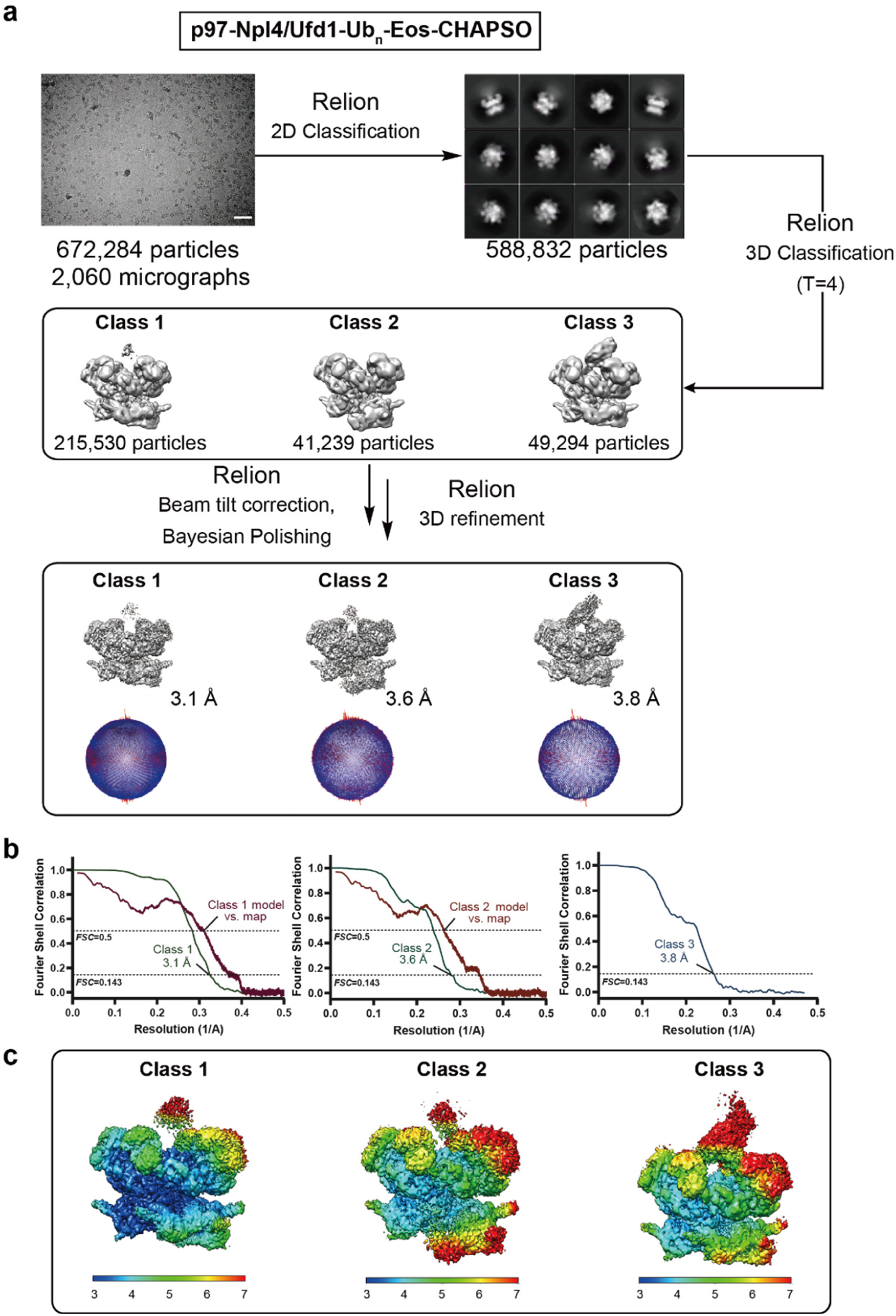
Single-particle cryo-EM analyses for the p97-Npl4/Ufd1-Ub_n_-Eos-CHAPSO dataset. **a,** The workflow of data processing. The dataset was subjected to particle selection, 2D classification, and multiple rounds of 3D classification. A representative micrograph (scale bar corresponds to 50 nm) and representative 2D class averages are shown. Three classes were resolved from the dataset, including class 1 (closed state), class 2 (open state), and class 3 (similar to the closed state but with the Npl4 density). The distributions of the Euler angles for each reconstruction are shown below the maps. **b,** Fourier shell correlation (FSC) curves of the masked maps after Relion postprocessing. The resolutions were determined by the FSC=0.143 criterion. The model vs. map FSC curves are also shown for class 1 and class 2 (red). **c,** Local resolutions of the maps calculated using Relion.

**Extended Data Fig. 3:**
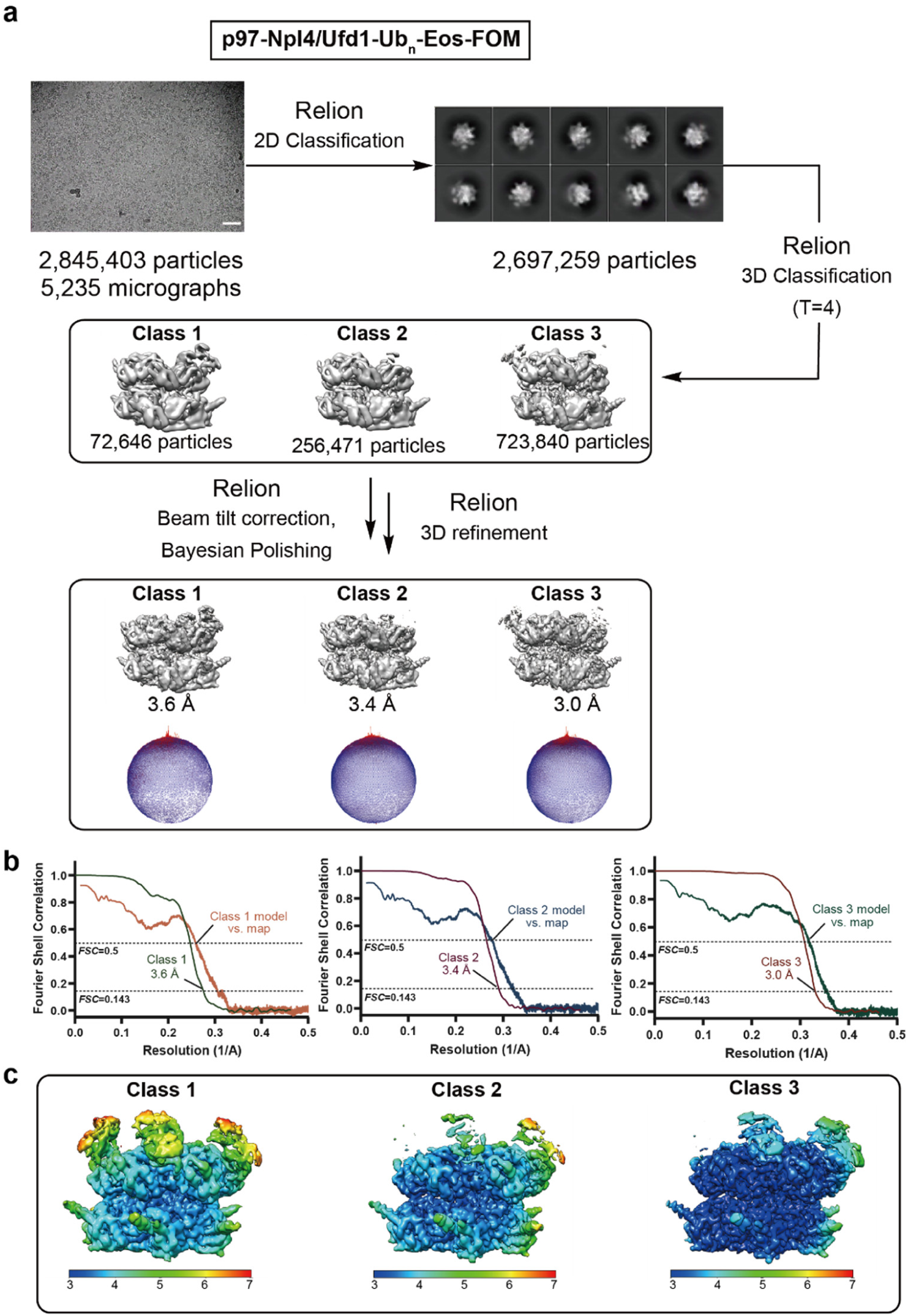
Single-particle cryo-EM analyses for the p97-Npl4/Ufd1-Ub_n_-Eos-FOM dataset. **a,** The workflow of data processing. The dataset was subjected to particle selection, 2D classification, and multiple rounds of 3D classification. A representative micrograph (scale bar corresponds to 50 nm) and representative 2D class averages are shown. Three classes were resolved from the dataset. The distributions of the Euler angles for each reconstruction are shown below the maps. **b,** FSC curves of the masked maps after Relion postprocessing. The resolutions were determined by the FSC=0.143 criterion. The model vs. map FSC curves for each class are also shown. **c,** Local resolutions of the maps calculated using Relion.

**Extended Data Fig. 4:**
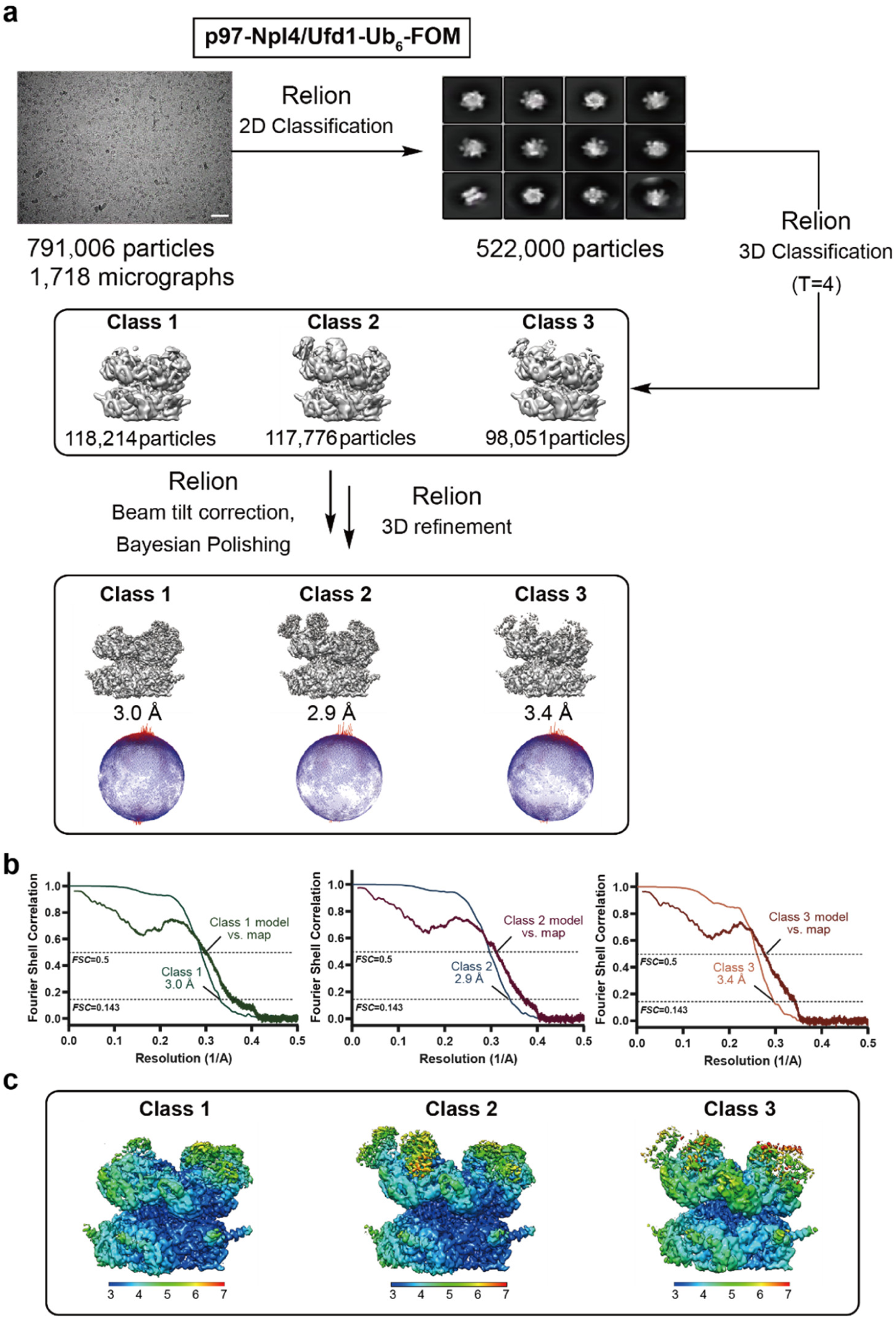
Single-particle cryo-EM analyses for the p97-Npl4/Ufd1-Ub_6_-FOM dataset. **a,** The workflow of data processing. The dataset was subjected to particle selection, 2D classification, and multiple rounds of 3D classification. A representative micrograph (scale bar corresponds to 50 nm) and representative 2D class averages are shown. Three classes were resolved from the dataset. The distributions of the Euler angles for each reconstruction are shown below the maps. **b,** FSC curves of the masked maps after Relion postprocessing. The resolutions were determined by the FSC=0.143 criterion. The model vs. map FSC curves for each class are also shown. **c,** Local resolutions of the maps calculated using Relion.

**Extended Data Fig. 5:**
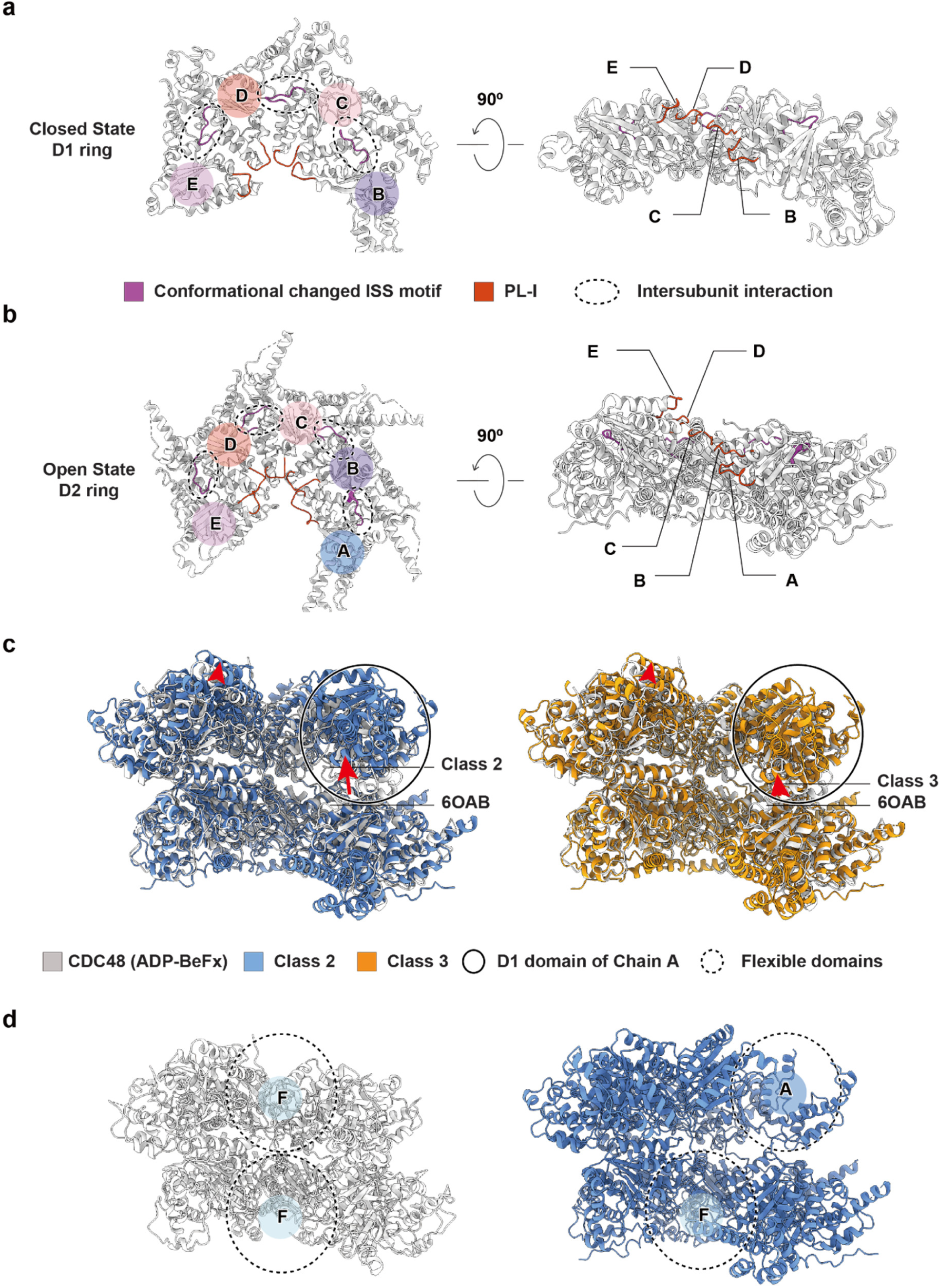
Comparisons of human p97 and yeast Cdc48 structures. **a-b,** Illustrations showing the arrangement of chains B-E in the D1 ring of the closed state (**a**) and chains A-E in the D2 ring of the open state (**b**). The labels of chains share the same color code as in Fig. 1E. **c,** Comparisons of cryo-EM models of translocating structures of p97 (class 2, medium blue; class 3, orange) from the p97-Npl4/Ufd1-Ub_n_-Eos-FOM dataset with Cdc48 (ADP-BeFx, PDB ID: 6OAB, light grey). The models were aligned based on the D2 ring. The relative shifts of the D1 domain in chain A are noted by red arrows. **d,** Illustrations showing the model of Cdc48 (ADP-BeFx, PDB ID: 6OAB, light grey, left panel) and the translocating structure of p97 (class 2, medium blue, right panel) without the flexible domains. The entire chain F in Cdc48 (ADP-BeFx) was flexible according to the previous publication ^19^. The D2 domain in chain F and D1 domain in chain A were flexible in this study. Both (positioned in dotted circles) were hid for comparison. Note that the staircases formed by D1 and D2 rings are in register for Cdc48 and out of register for p97.

**Extended Data Fig. 6:**
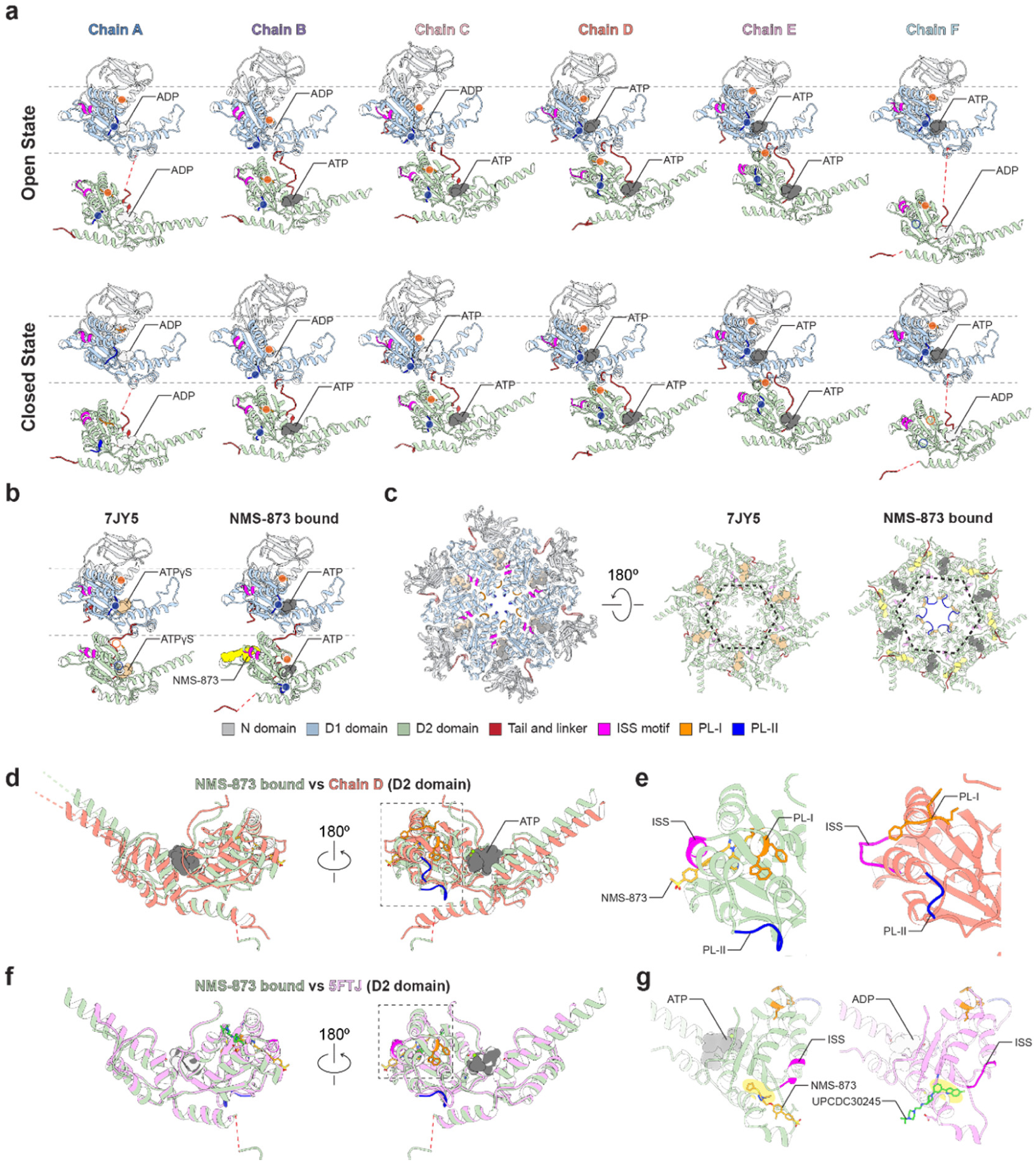
Comparison of various p97 structures. **a,** Unrolling of the open (top row) and closed (bottom row) states of p97. The lateral and vertical movement of D2 domains can be visualized. The positions of pore loops are marked with solid dots. Unresolved pore loops are marked by open circles. **b,** Unrolling of the ATPγS-bound nontranslocating structure (PDB ID: 7JY5) and NMS-873-bound structure. Only one chain of each structure is shown since both structures are sixfold symmetric. **c,** A superimposition of 7JY5- and NMS-873-bound structures. Left: a top view of the N and D1 domains; Right: a bottom view of the D2 ring. The opening of the D2 ring is marked by a dotted hexagon. Panels **a, b,** and **c** share the same color code. **d,** A comparison of the D2 domain in the NMS-873-bound structure and that in chain D of the open state. The superimposition is based on the α/β subdomain. **e,** Magnified views of the dotted box in panel **d**, showing the conformations of the PL-I, PL-II, and ISS motifs. **f,** A comparison of the D2 domain in NMS-873- and UPCDC30245 (PDB ID: 5FTJ)-bound structures. The superimposition is based on the α/β subdomain. **g,** Magnified views of the dotted box in panel **f**, showing the binding sites of the compounds and the conformations of the ISS motif.

**Extended Data Fig. 7:**
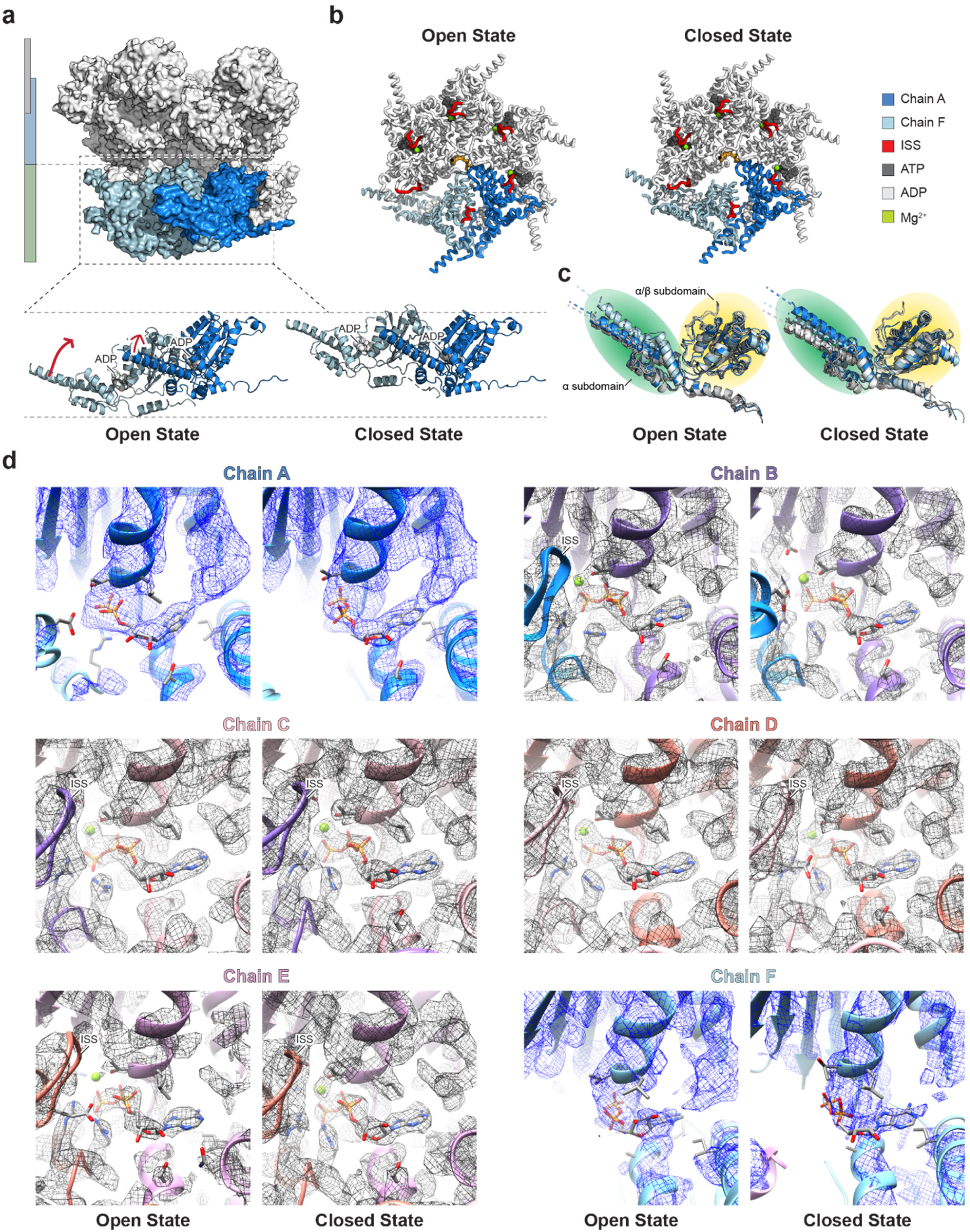
The power stroke motion and the nucleotide binding sites of the D2 domains. **a,** A superimposition of the open and closed state structures, with two D2 domains (chain A and chain F) magnified. The relative positions of the D2 domains in chain A and chain F can be visualized. **b,** Top views of the D2 ring in the open and closed states, highlighting chain A, chain F, and the ISS motifs. **c,** Superimpositions of all D2 domains in the open and closed states based on the α/β subdomain. The directions of α7 helices are highlighted by dotted lines for comparison (dark blue, chain A; light blue, chain F; and gray, other chains). **d,** Individual nucleotide binding sites of the D2 domains in the open and closed states. Blue mesh: unsharpened map at contour level; Gray mesh: sharpened map at contour level.;

**Extended Data Fig. 8:**
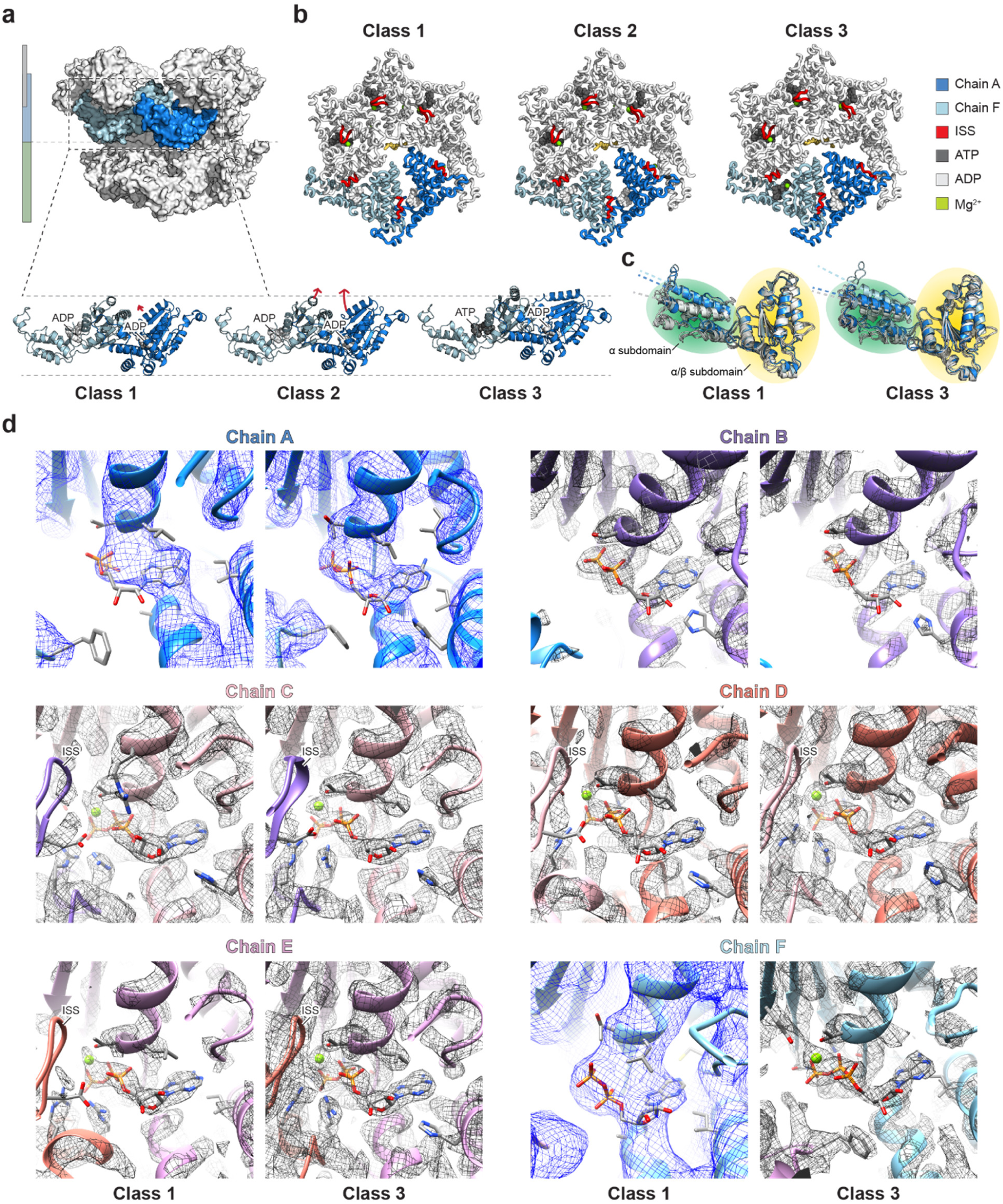
The power stroke motion and the nucleotide binding sites of the D1 domains. **a,** A superimposition of three structures (classes 1, 2, and 3) from the p97-Npl4/Ufd1-Ub_n_-Eos-FOM dataset, with two D1 domains (chain A and chain F) magnified. The relative positions of the D1 domains in chain A and chain F can be visualized. **b,** Top views of the D1 ring in the three classes, highlighting chain A, chain F, and the ISS motifs. **c,** Superimpositions of all D1 domains in class 1 and class 3 based on the α/β subdomain. The directions of α7 helices are highlighted by dotted lines for comparison (dark blue, chain A; light blue, chain F; and gray, other chains). **d,** Individual nucleotide binding sites of the D1 domains in class 1 and class 3. Blue mesh: Unsharpened map at contour level; Gray mesh: sharpened map at contour level.;

**Extended Data Fig. 9:**
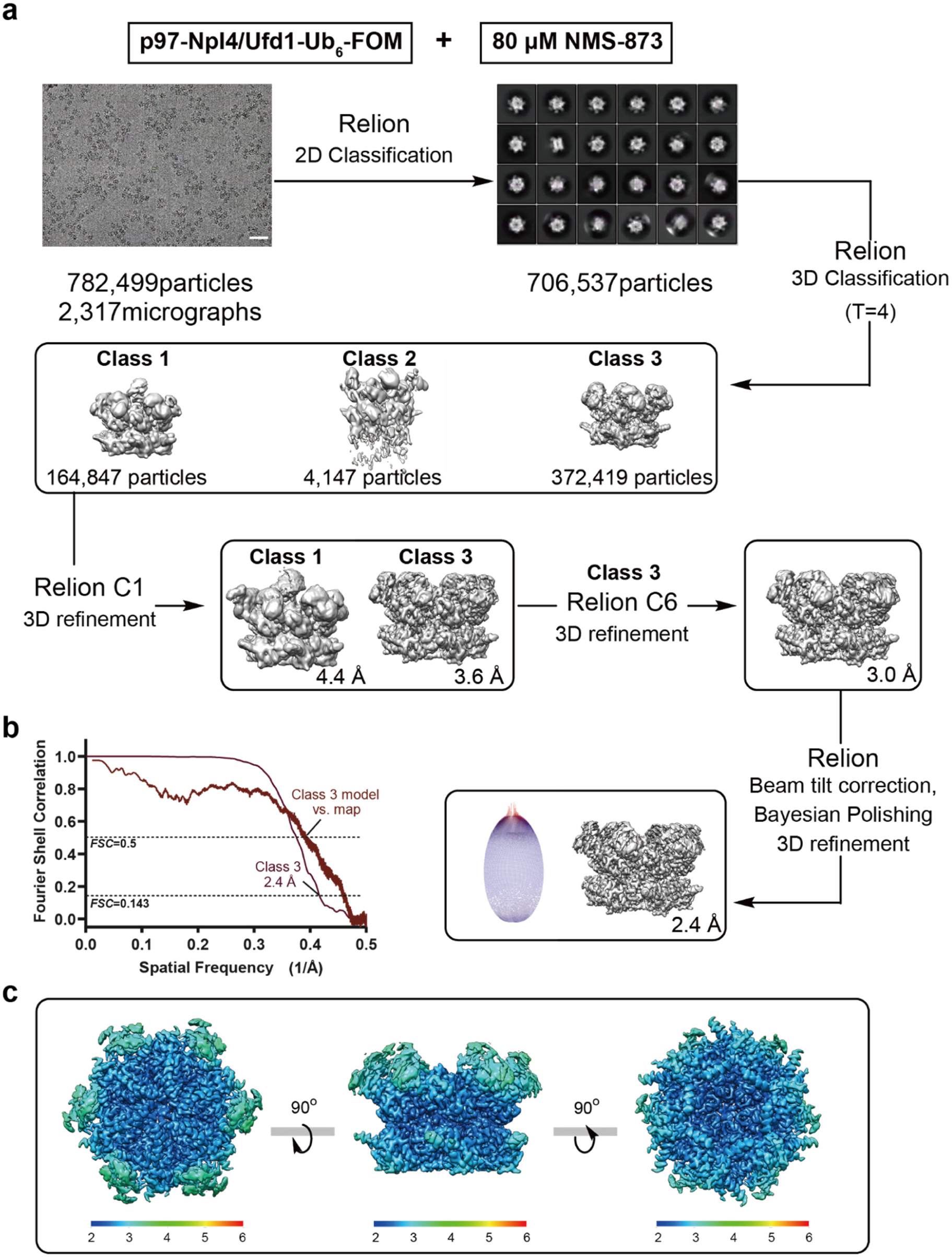
Single-particle cryo-EM analyses for the p97-Npl4/Ufd1-Ub_6_ complex in the presence of 80 μM NMS-873. **a,** The workflow of data processing. The dataset was subjected to particle selection, 2D classification, and multiple rounds of 3D classification. A representative micrograph (scale bar corresponds to 50 nm) and representative 2D class averages are shown. A single class was resolved from the dataset. The distribution of the Euler angles is shown next to the map. **b,** The FSC curve of the masked map after Relion postprocessing. The resolution was determined by the FSC=0.143 criterion. The model vs. map FSC curve is also shown. **c,** Local resolution of the map calculated using Relion.

**Extended Data Fig. 10:**
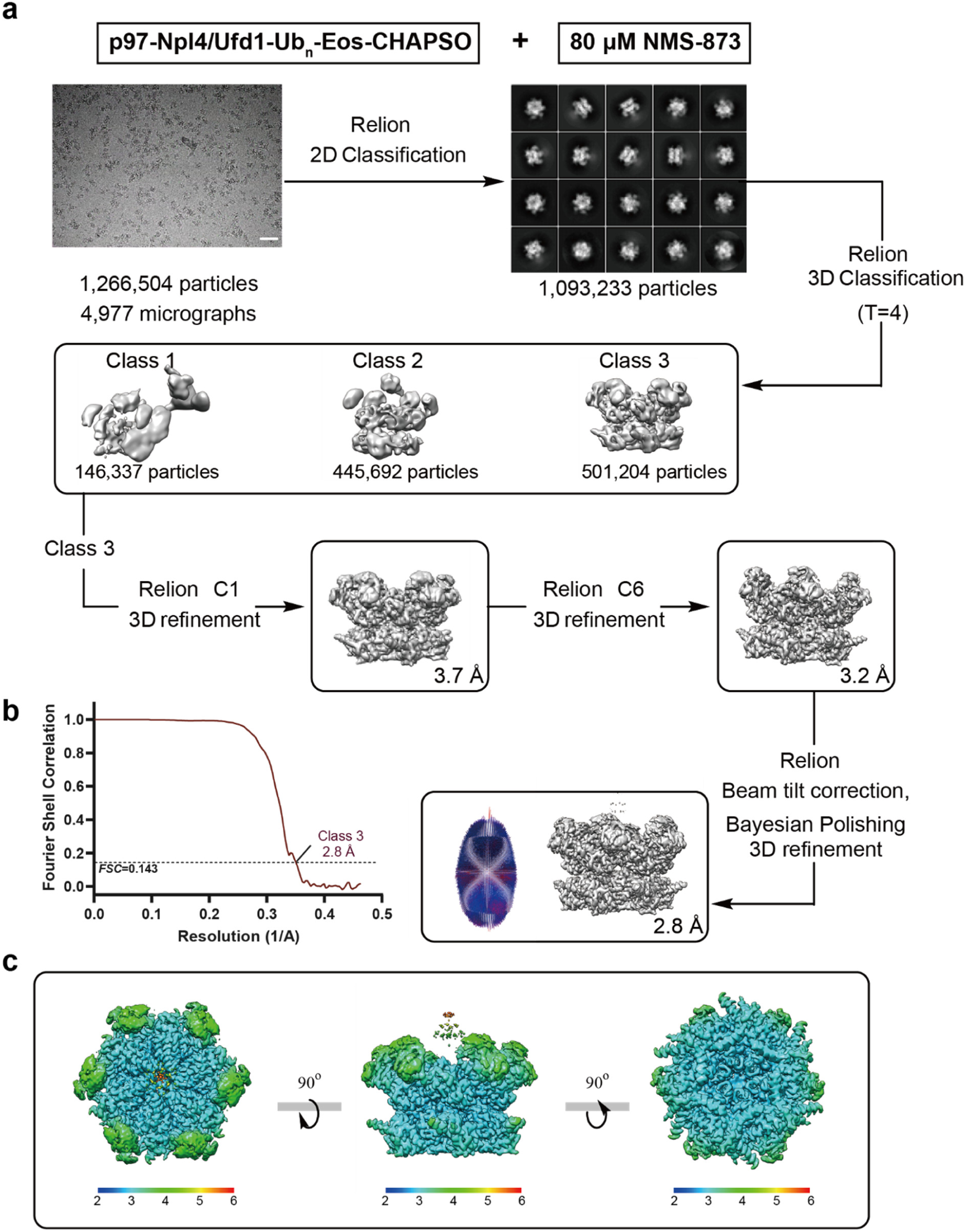
Single-particle cryo-EM analyses for the p97-Npl4/Ufd1-Ub_n_-Eos complex in the presence of 80 μM NMS-873. **a,** The workflow of data processing. The dataset was subjected to particle selection, 2D classification, and multiple rounds of 3D classification. A representative micrograph (scale bar corresponds to 50 nm) and representative 2D class averages are shown. A single class was resolved from the dataset. The distribution of the Euler angles is shown next to the map. **b,** The FSC curve of the masked map after Relion postprocessing. The resolution was determined by the FSC=0.143 criterion. **c,** Local resolution of the map calculated using Relion.

**Extended Data Fig. 11:**
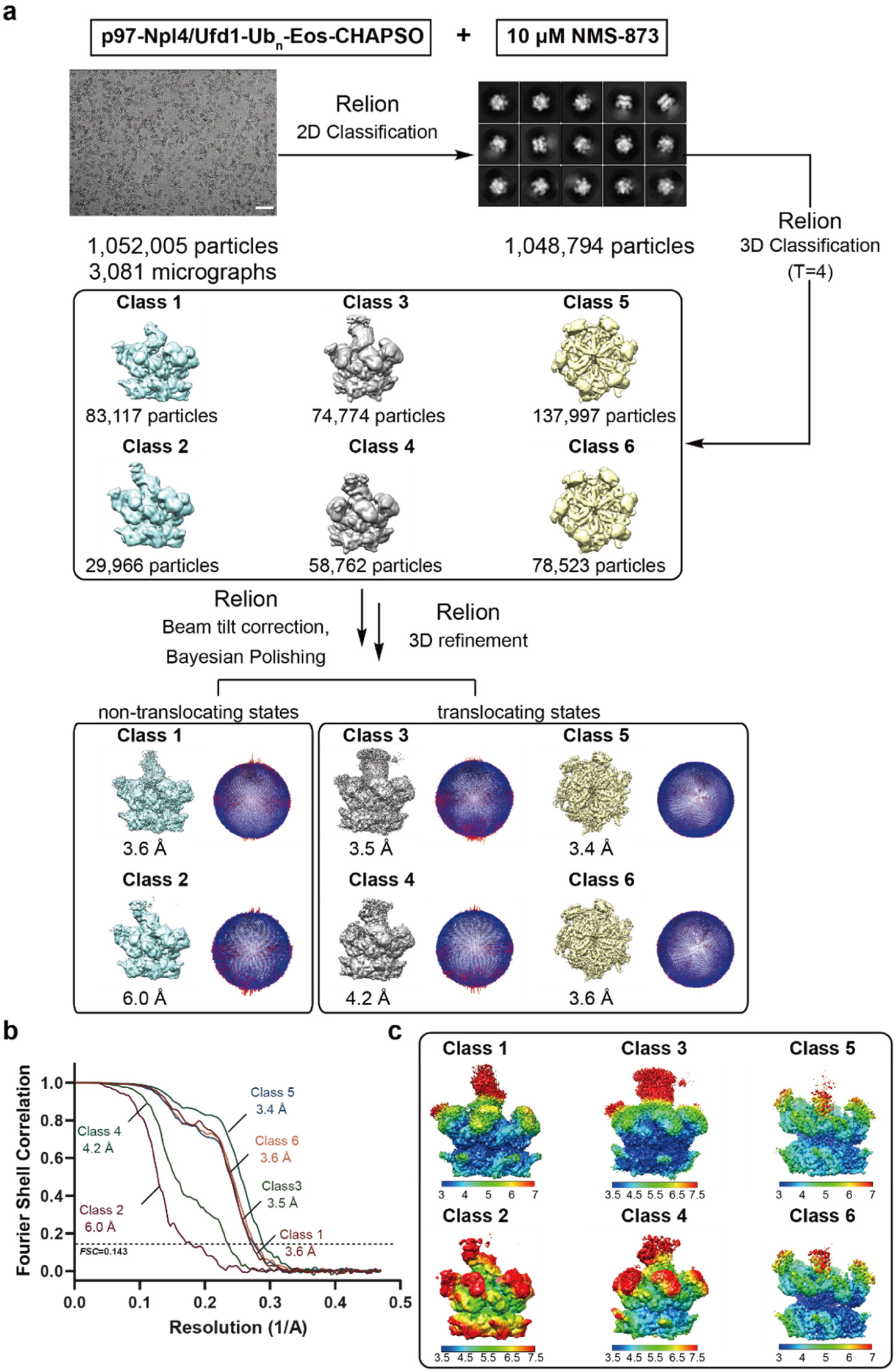
Single-particle cryo-EM analyses for the p97-Npl4/Ufd1-Ub_n_-Eos complex in the presence of 10 μM NMS-873. **a,** The workflow of data processing. The dataset was subjected to particle selection, 2D classification, and multiple rounds of 3D classification. A representative micrograph (scale bar corresponds to 50 nm) and representative 2D class averages are shown. Six classes were resolved from the dataset, including two in nontranslocating states (Class 1 and Class 2) and four in translocating states (Class 3 through Class 6). The distributions of the Euler angles for each reconstruction are shown next to the maps. **b,** FSC curves of the masked maps after Relion postprocessing. The resolutions were determined by the FSC=0.143 criterion. **c,** Local resolutions of the maps calculated using Relion.

**Extended Data Table 1:**
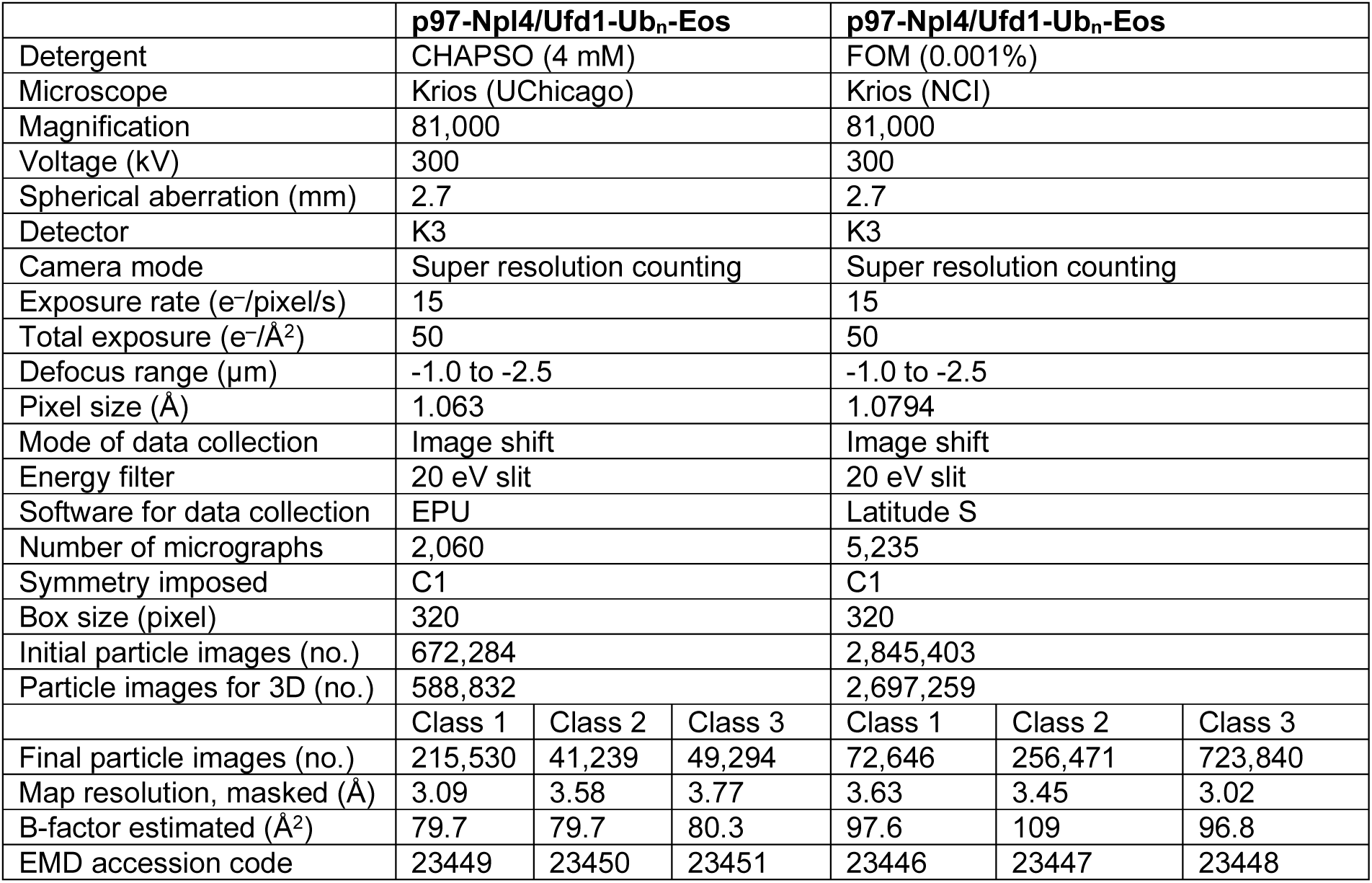
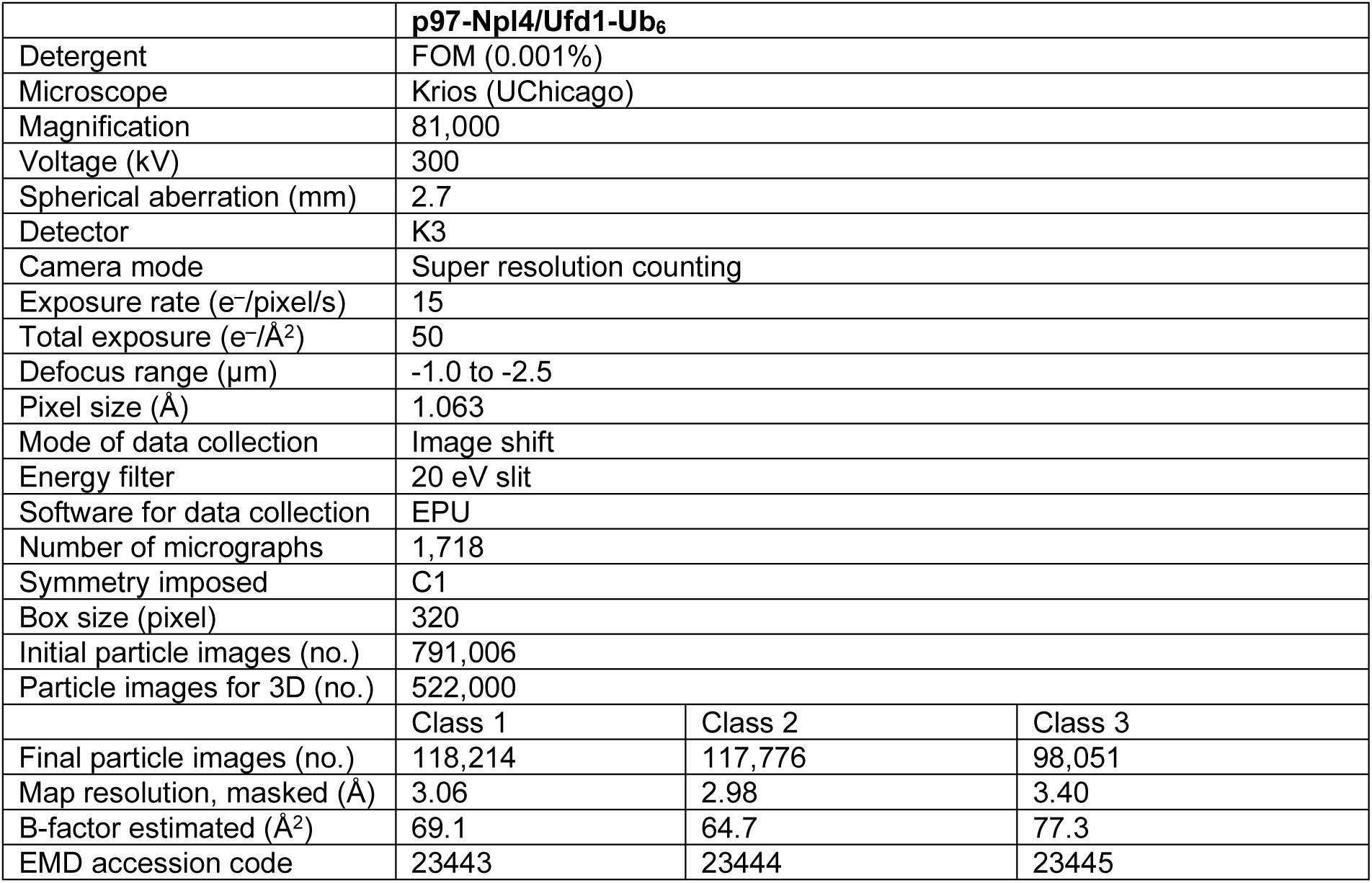
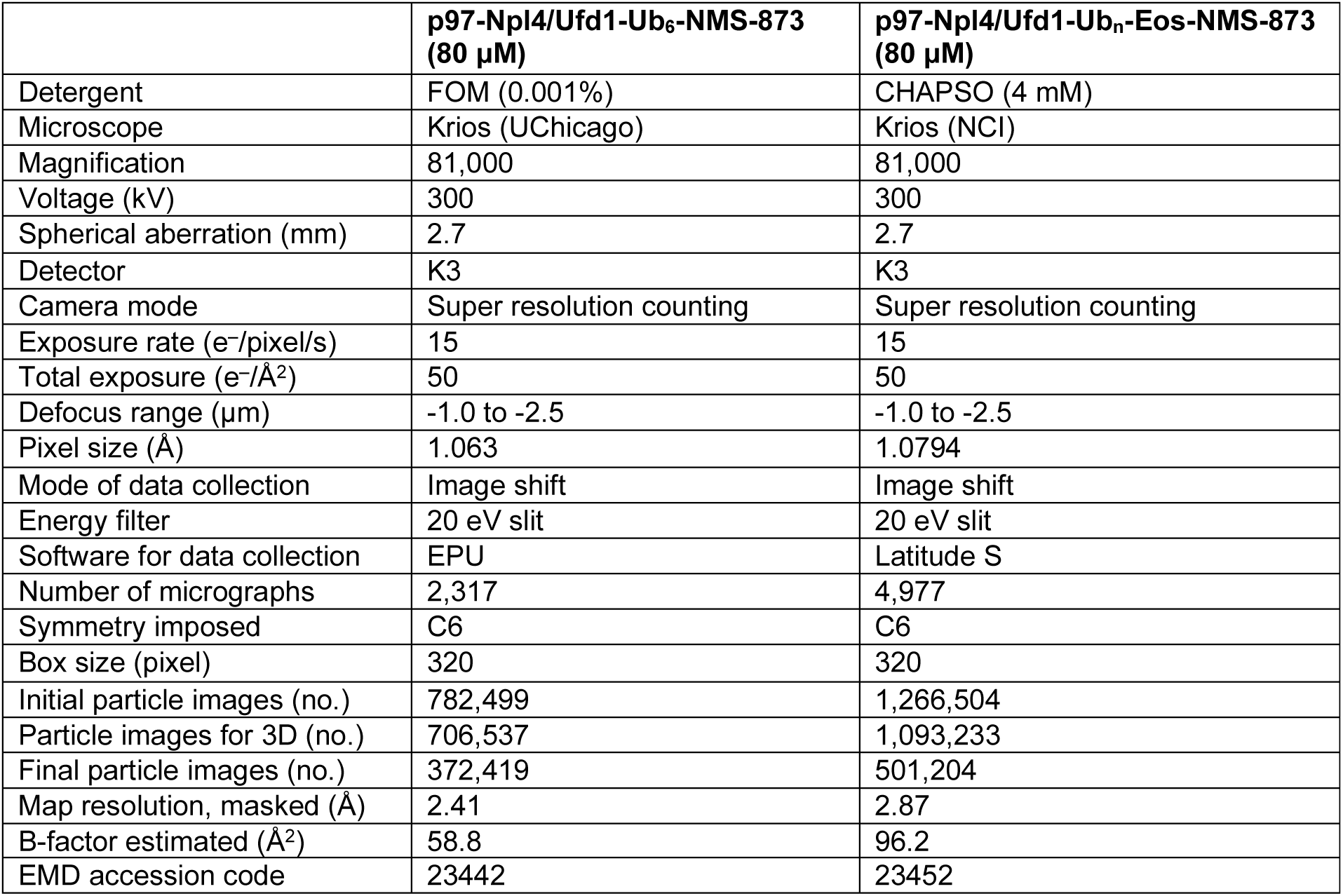
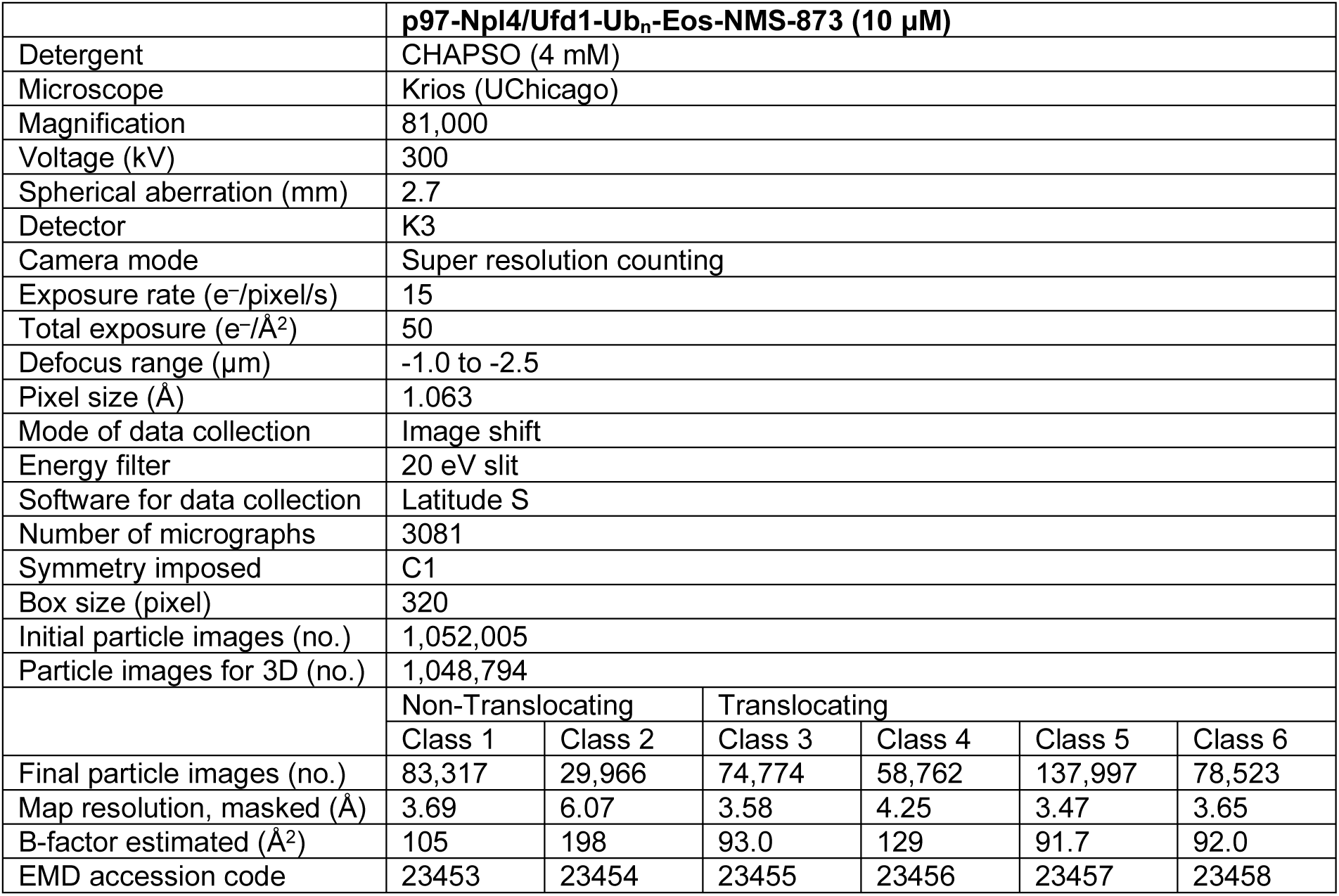
Statistics of cryo-EM data collection and processing

|  | **p97-Npl4/Ufd1-Ub<sub>n</sub>-Eos** |  |  | **p97-Npl4/Ufd1-Ub<sub>n</sub>-Eos** |  |  |
| --- | --- | --- | --- | --- | --- | --- |
| Detergent | CHAPSO (4 mM) |  |  | FOM (0.001%) |  |  |
| Microscope | Krios (UChicago) |  |  | Krios (NCI) |  |  |
| Magnification | 81,000 |  |  | 81,000 |  |  |
| Voltage (kV) | 300 |  |  | 300 |  |  |
| Spherical aberration (mm) | 2.7 |  |  | 2.7 |  |  |
| Detector | K3 |  |  | K3 |  |  |
| Camera mode | Super resolution counting |  |  | Super resolution counting |  |  |
| Exposure rate (e<sup>-</sup>/pixel/s) | 15 |  |  | 15 |  |  |
| Total exposure (e<sup>-</sup>/Å<sup>2</sup>) | 50 |  |  | 50 |  |  |
| Defocus range (μm) | -1.0 to -2.5 |  |  | -1.0 to -2.5 |  |  |
| Pixel size (Å) | 1.063 |  |  | 1.0794 |  |  |
| Mode of data collection | Image shift |  |  | Image shift |  |  |
| Energy filter | 20 eV slit |  |  | 20 eV slit |  |  |
| Software for data collection | EPU |  |  | Latitude S |  |  |
| Number of micrographs | 2,060 |  |  | 5,235 |  |  |
| Symmetry imposed | C1 |  |  | C1 |  |  |
| Box size (pixel) | 320 |  |  | 320 |  |  |
| Initial particle images (no.) | 672,284 |  |  | 2,845,403 |  |  |
| Particle images for 3D (no.) | 588,832 |  |  | 2,697,259 |  |  |
|  | Class 1 | Class 2 | Class 3 | Class 1 | Class 2 | Class 3 |
| Final particle images (no.) | 215,530 | 41,239 | 49,294 | 72,646 | 256,471 | 723,840 |
| Map resolution, masked (Å) | 3.09 | 3.58 | 3.77 | 3.63 | 3.45 | 3.02 |
| B-factor estimated (Å<sup>2</sup>) | 79.7 | 79.7 | 80.3 | 97.6 | 109 | 96.8 |
| EMD accession code | 23449 | 23450 | 23451 | 23446 | 23447 | 23448 |

|  |  |  |  |
| --- | --- | --- | --- |
|  | **p97-Npl4/Ufd1-Ub<sub>6</sub>** |  |  |
| Detergent | FOM (0.001%) |  |  |
| Microscope | Krios (UChicago) |  |  |
| Magnification | 81,000 |  |  |
| Voltage (kV) | 300 |  |  |
| Spherical aberration (mm) | 2.7 |  |  |
| Detector | K3 |  |  |
| Camera mode | Super resolution counting |  |  |
| Exposure rate (e<sup>-</sup>/pixel/s) | 15 |  |  |
| Total exposure (e<sup>-</sup>/Å<sup>2</sup>) | 50 |  |  |
| Defocus range (μm) | -1.0 to -2.5 |  |  |
| Pixel size (Å) | 1.063 |  |  |
| Mode of data collection | Image shift |  |  |
| Energy filter | 20 eV slit |  |  |
| Software for data collection | EPU |  |  |
| Number of micrographs | 1,718 |  |  |
| Symmetry imposed | C1 |  |  |
| Box size (pixel) | 320 |  |  |
| Initial particle images (no.) | 791,006 |  |  |
| Particle images for 3D (no.) | 522,000 |  |  |
|  | Class 1 | Class 2 | Class 3 |
| Final particle images (no.) | 118,214 | 117,776 | 98,051 |
| Map resolution, masked (Å) | 3.06 | 2.98 | 3.40 |
| B-factor estimated (Å<sup>2</sup>) | 69.1 | 64.7 | 77.3 |
| EMD accession code | 23443 | 23444 | 23445 |

|  | **p97-Npl4/Ufd1-Ub<sub>6</sub>-NMS-873**<br>**(80 μM)** | **p97-Npl4/Ufd1-Ub<sub>n</sub>-Eos-NMS-873**<br>**(80 μM)** |
| --- | --- | --- |
| Detergent | FOM (0.001%) | CHAPSO (4 mM) |
| Microscope | Krios (UChicago) | Krios (NCI) |
| Magnification | 81,000 | 81,000 |
| Voltage (kV) | 300 | 300 |
| Spherical aberration (mm) | 2.7 | 2.7 |
| Detector | K3 | K3 |
| Camera mode | Super resolution counting | Super resolution counting |
| Exposure rate (e<sup>-</sup>/pixel/s) | 15 | 15 |
| Total exposure (e<sup>-</sup>/Å<sup>2</sup>) | 50 | 50 |
| Defocus range (μm) | -1.0 to -2.5 | -1.0 to -2.5 |
| Pixel size (Å) | 1.063 | 1.0794 |
| Mode of data collection | Image shift | Image shift |
| Energy filter | 20 eV slit | 20 eV slit |
| Software for data collection | EPU | Latitude S |
| Number of micrographs | 2,317 | 4,977 |
| Symmetry imposed | C6 | C6 |
| Box size (pixel) | 320 | 320 |
| Initial particle images (no.) | 782,499 | 1,266,504 |
| Particle images for 3D (no.) | 706,537 | 1,093,233 |
| Final particle images (no.) | 372,419 | 501,204 |
| Map resolution, masked (Å) | 2.41 | 2.87 |
| B-factor estimated (Å<sup>2</sup>) | 58.8 | 96.2 |
| EMD accession code | 23442 | 23452 |
|  | **p97-Npl4/Ufd1-Ub<sub>n</sub>-Eos-NMS-873 (10 µM)** |  |  |  |  |  |
| Detergent | CHAPSO (4 mM) |  |  |  |  |  |
| Microscope | Krios (UChicago) |  |  |  |  |  |
| Magnification | 81,000 |  |  |  |  |  |
| Voltage (kV) | 300 |  |  |  |  |  |
| Spherical aberration (mm) | 2.7 |  |  |  |  |  |
| Detector | K3 |  |  |  |  |  |
| Camera mode | Super resolution counting |  |  |  |  |  |
| Exposure rate (e<sup>-</sup>/pixel/s) | 15 |  |  |  |  |  |
| Total exposure (e<sup>-</sup>/Å<sup>2</sup>) | 50 |  |  |  |  |  |
| Defocus range (µm) | -1.0 to -2.5 |  |  |  |  |  |
| Pixel size (Å) | 1.063 |  |  |  |  |  |
| Mode of data collection | Image shift |  |  |  |  |  |
| Energy filter | 20 eV slit |  |  |  |  |  |
| Software for data collection | Latitude S |  |  |  |  |  |
| Number of micrographs | 3081 |  |  |  |  |  |
| Symmetry imposed | C1 |  |  |  |  |  |
| Box size (pixel) | 320 |  |  |  |  |  |
| Initial particle images (no.) | 1,052,005 |  |  |  |  |  |
| Particle images for 3D (no.) | 1,048,794 |  |  |  |  |  |
|  | Non-Translocating |  | Translocating |  |  |  |
|  | Class 1 | Class 2 | Class 3 | Class 4 | Class 5 | Class 6 |
| Final particle images (no.) | 83,317 | 29,966 | 74,774 | 58,762 | 137,997 | 78,523 |
| Map resolution, masked (Å) | 3.69 | 6.07 | 3.58 | 4.25 | 3.47 | 3.65 |
| B-factor estimated (Å<sup>2</sup>) | 105 | 198 | 93.0 | 129 | 91.7 | 92.0 |
| EMD accession code | 23453 | 23454 | 23455 | 23456 | 23457 | 23458 |

**Extended Data Table 2:**
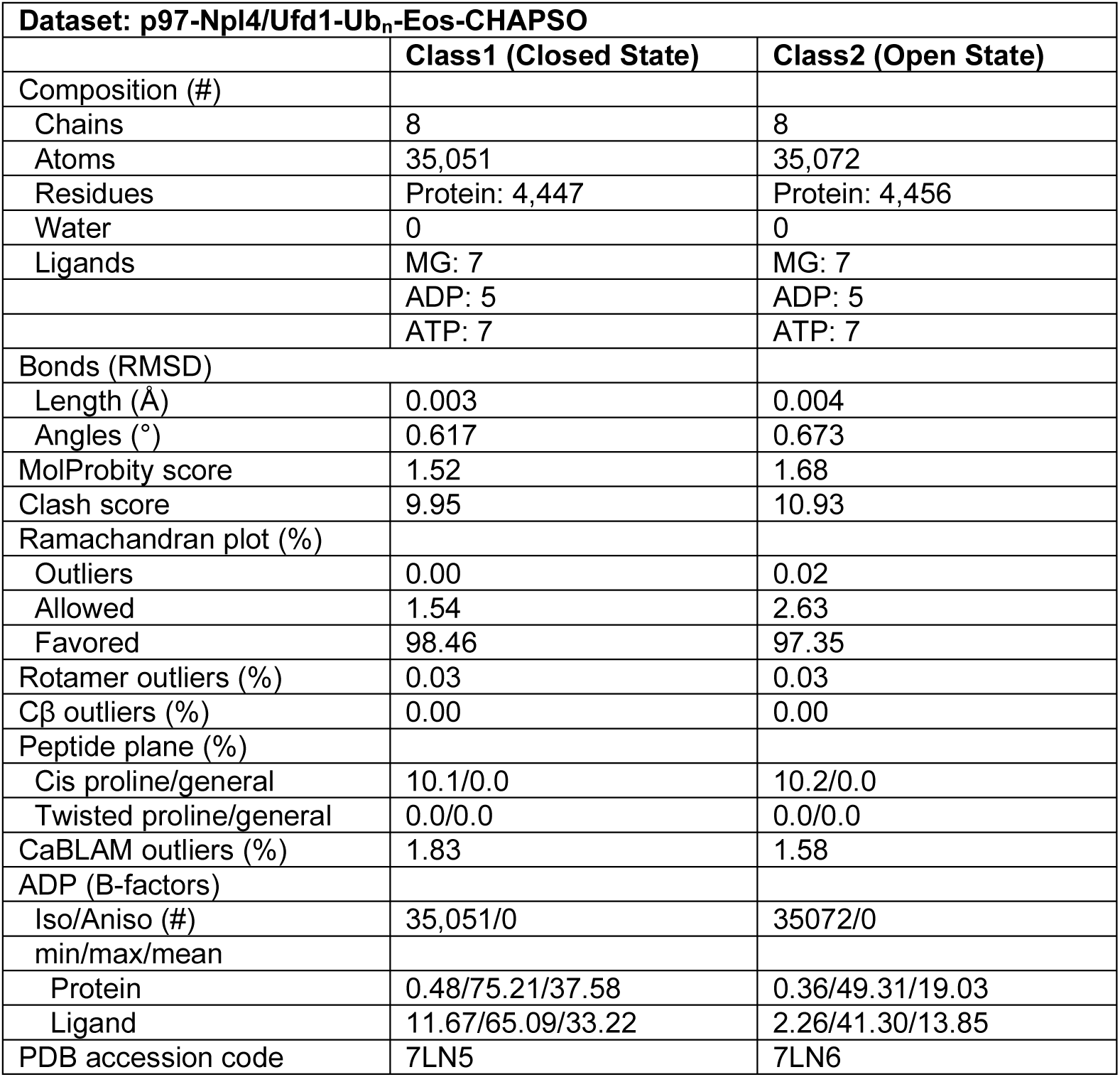
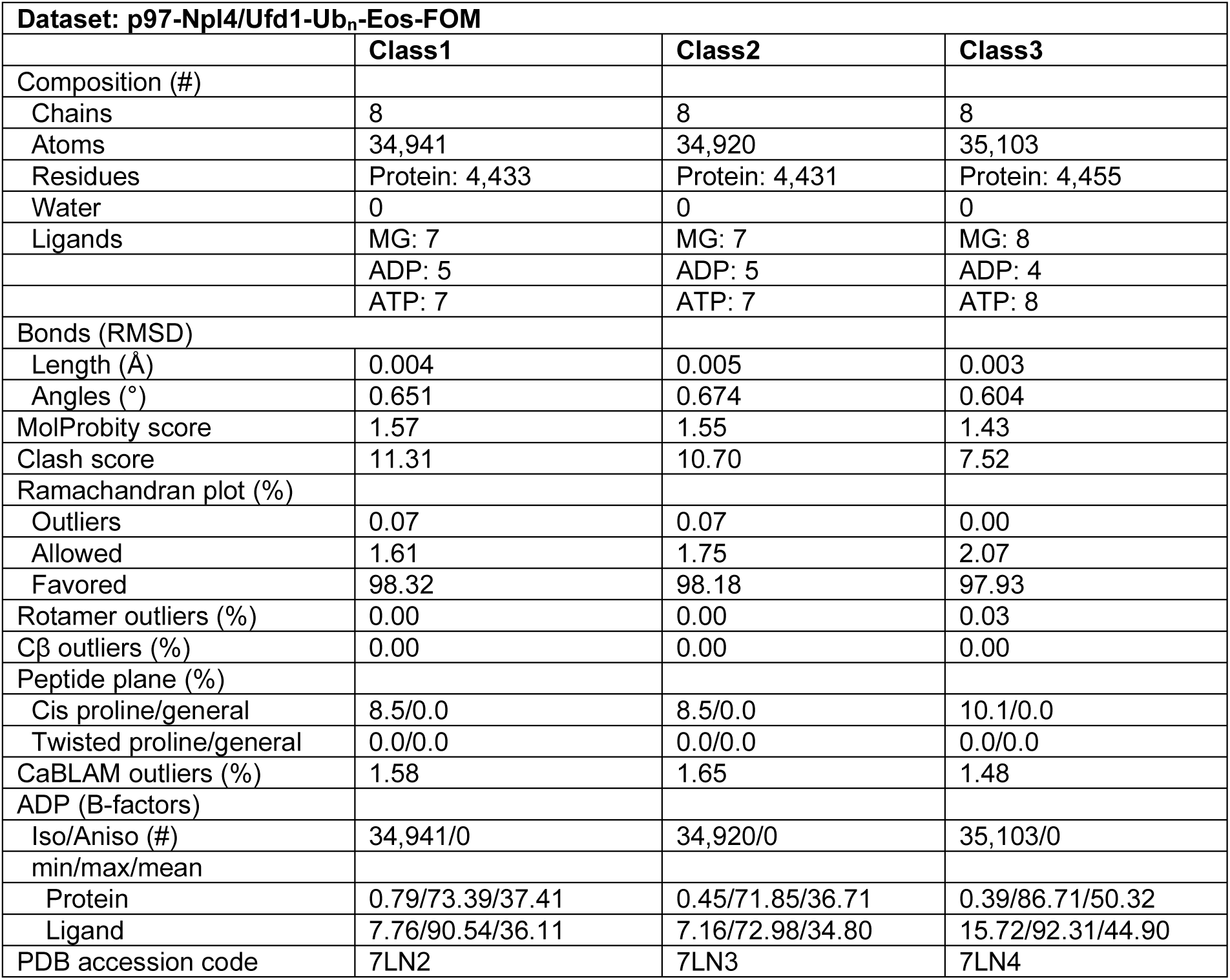
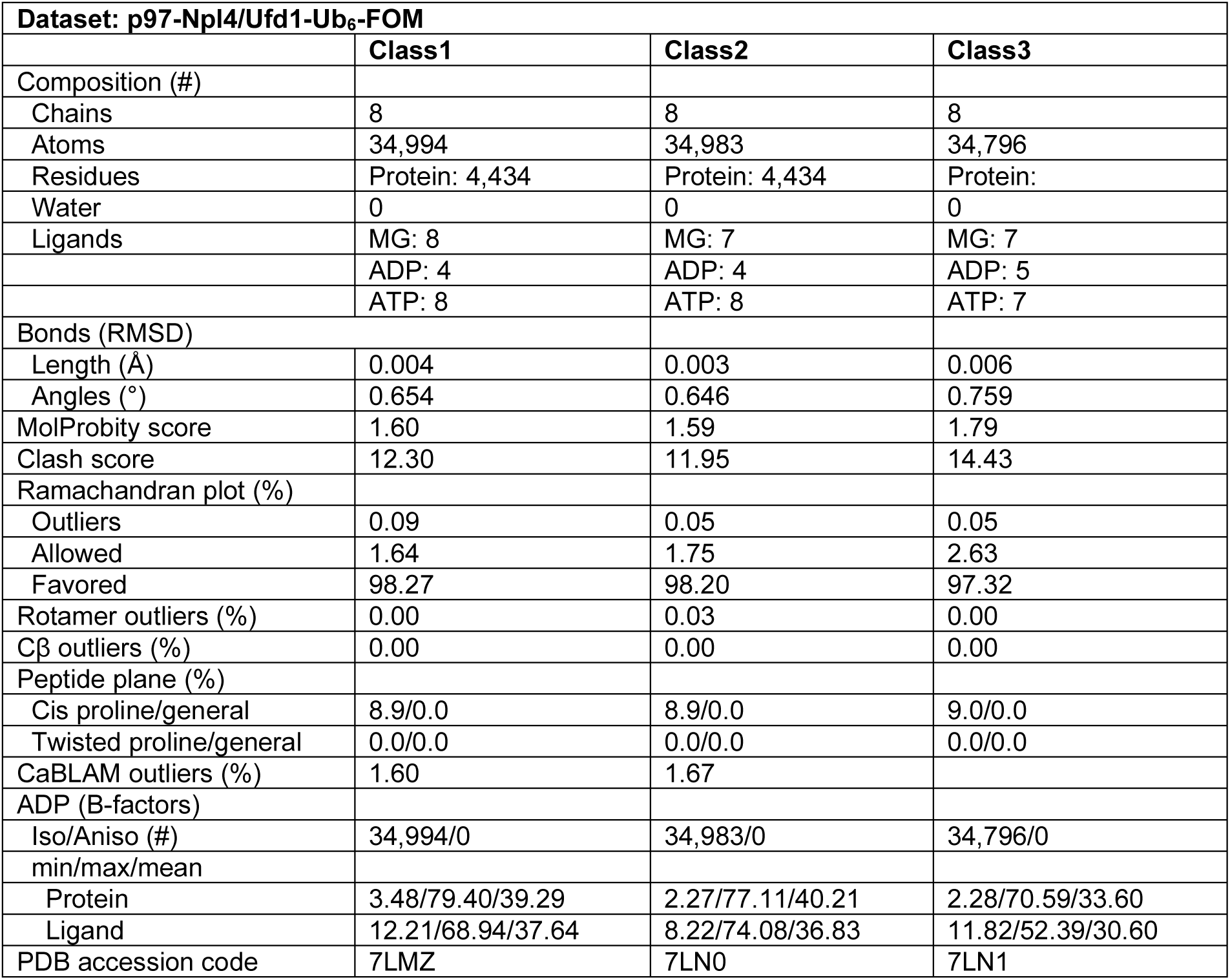
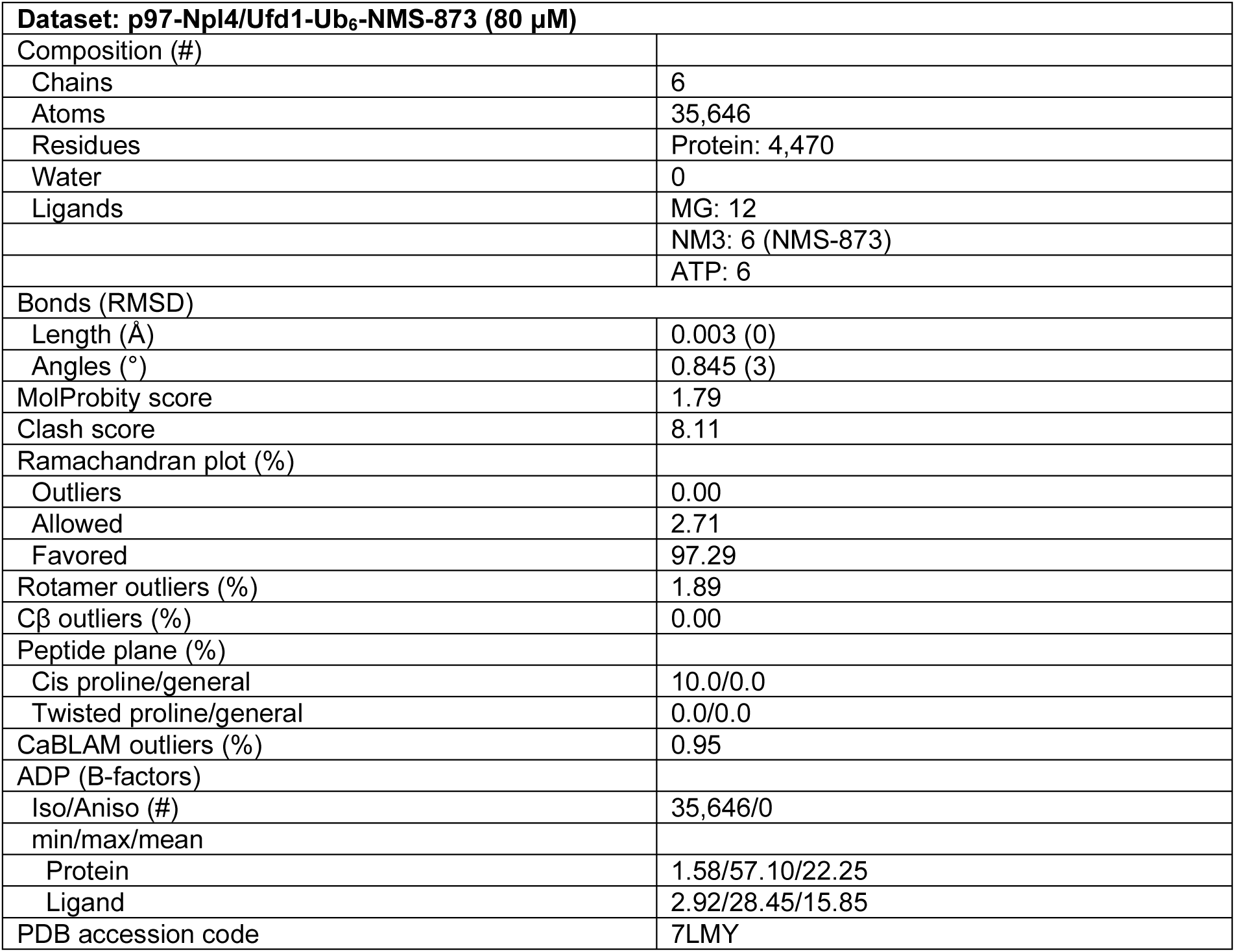
Statistics of cryo-EM model refinement and geometry

| **Dataset: p97-Npl4/Ufd1-Ub<sub>n</sub>-Eos-CHAPSO** |  |  |
| --- | --- | --- |
|  | **Class1 (Closed State)** | **Class2 (Open State)** |
| Composition (#) |  |  |
| Chains | 8 | 8 |
| Atoms | 35,051 | 35,072 |
| Residues | Protein: 4,447 | Protein: 4,456 |
| Water | 0 | 0 |
| Ligands | MG: 7 | MG: 7 |
|  | ADP: 5 | ADP: 5 |
|  | ATP: 7 | ATP: 7 |
| Bonds (RMSD) |  |  |
| Length (Å) | 0.003 | 0.004 |
| Angles (°) | 0.617 | 0.673 |
| MolProbity score | 1.52 | 1.68 |
| Clash score | 9.95 | 10.93 |
| Ramachandran plot (%) |  |  |
| Outliers | 0.00 | 0.02 |
| Allowed | 1.54 | 2.63 |
| Favored | 98.46 | 97.35 |
| Rotamer outliers (%) | 0.03 | 0.03 |
| Cβ outliers (%) | 0.00 | 0.00 |
| Peptide plane (%) |  |  |
| Cis proline/general | 10.1/0.0 | 10.2/0.0 |
| Twisted proline/general | 0.0/0.0 | 0.0/0.0 |
| CaBLAM outliers (%) | 1.83 | 1.58 |
| ADP (B-factors) |  |  |
| Iso/Aniso (#) | 35,051/0 | 35072/0 |
| min/max/mean |  |  |
| Protein | 0.48/75.21/37.58 | 0.36/49.31/19.03 |
| Ligand | 11.67/65.09/33.22 | 2.26/41.30/13.85 |
| PDB accession code | 7LN5 | 7LN6 |

| Dataset: p97-Npl4/Ufd1-Ub<sub>n</sub>-Eos-FOM |  |  |  |
| --- | --- | --- | --- |
|  | Class1 | Class2 | Class3 |
| Composition (#) |  |  |  |
| Chains | 8 | 8 | 8 |
| Atoms | 34,941 | 34,920 | 35,103 |
| Residues | Protein: 4,433 | Protein: 4,431 | Protein: 4,455 |
| Water | 0 | 0 | 0 |
| Ligands | MG: 7 | MG: 7 | MG: 8 |
|  | ADP: 5 | ADP: 5 | ADP: 4 |
|  | ATP: 7 | ATP: 7 | ATP: 8 |
| Bonds (RMSD) |  |  |  |
| Length (Å) | 0.004 | 0.005 | 0.003 |
| Angles (°) | 0.651 | 0.674 | 0.604 |
| MolProbity score | 1.57 | 1.55 | 1.43 |
| Clash score | 11.31 | 10.70 | 7.52 |
| Ramachandran plot (%) |  |  |  |
| Outliers | 0.07 | 0.07 | 0.00 |
| Allowed | 1.61 | 1.75 | 2.07 |
| Favored | 98.32 | 98.18 | 97.93 |
| Rotamer outliers (%) | 0.00 | 0.00 | 0.03 |
| Cβ outliers (%) | 0.00 | 0.00 | 0.00 |
| Peptide plane (%) |  |  |  |
| Cis proline/general | 8.5/0.0 | 8.5/0.0 | 10.1/0.0 |
| Twisted proline/general | 0.0/0.0 | 0.0/0.0 | 0.0/0.0 |
| CaBLAM outliers (%) | 1.58 | 1.65 | 1.48 |
| ADP (B-factors) |  |  |  |
| Iso/Aniso (#) | 34,941/0 | 34,920/0 | 35,103/0 |
| min/max/mean |  |  |  |
| Protein | 0.79/73.39/37.41 | 0.45/71.85/36.71 | 0.39/86.71/50.32 |
| Ligand | 7.76/90.54/36.11 | 7.16/72.98/34.80 | 15.72/92.31/44.90 |
| PDB accession code | 7LN2 | 7LN3 | 7LN4 |

| Dataset: p97-Npl4/Ufd1-Ub<sub>6</sub>-FOM |  |  |  |
| --- | --- | --- | --- |
|  | Class1 | Class2 | Class3 |
| Composition (#) |  |  |  |
| Chains | 8 | 8 | 8 |
| Atoms | 34,994 | 34,983 | 34,796 |
| Residues | Protein: 4,434 | Protein: 4,434 | Protein: |
| Water | 0 | 0 | 0 |
| Ligands | MG: 8 | MG: 7 | MG: 7 |
|  | ADP: 4 | ADP: 4 | ADP: 5 |
|  | ATP: 8 | ATP: 8 | ATP: 7 |
| Bonds (RMSD) |  |  |  |
| Length (Å) | 0.004 | 0.003 | 0.006 |
| Angles (°) | 0.654 | 0.646 | 0.759 |
| MolProbity score | 1.60 | 1.59 | 1.79 |
| Clash score | 12.30 | 11.95 | 14.43 |
| Ramachandran plot (%) |  |  |  |
| Outliers | 0.09 | 0.05 | 0.05 |
| Allowed | 1.64 | 1.75 | 2.63 |
| Favored | 98.27 | 98.20 | 97.32 |
| Rotamer outliers (%) | 0.00 | 0.03 | 0.00 |
| Cβ outliers (%) | 0.00 | 0.00 | 0.00 |
| Peptide plane (%) |  |  |  |
| Cis proline/general | 8.9/0.0 | 8.9/0.0 | 9.0/0.0 |
| Twisted proline/general | 0.0/0.0 | 0.0/0.0 | 0.0/0.0 |
| CaBLAM outliers (%) | 1.60 | 1.67 |  |
| ADP (B-factors) |  |  |  |
| Iso/Aniso (#) | 34,994/0 | 34,983/0 | 34,796/0 |
| min/max/mean |  |  |  |
| Protein | 3.48/79.40/39.29 | 2.27/77.11/40.21 | 2.28/70.59/33.60 |
| Ligand | 12.21/68.94/37.64 | 8.22/74.08/36.83 | 11.82/52.39/30.60 |
| PDB accession code | 7LMZ | 7LN0 | 7LN1 |

|  |  |
| --- | --- |
| **Dataset: p97-Npl4/Ufd1-Ub<sub>6</sub>-NMS-873 (80 µM)** |  |
| Composition (#) |  |
| Chains | 6 |
| Atoms | 35,646 |
| Residues | Protein: 4,470 |
| Water | 0 |
| Ligands | MG: 12 |
|  | NM3: 6 (NMS-873) |
|  | ATP: 6 |
| Bonds (RMSD) |  |
| Length (Å) | 0.003 (0) |
| Angles (°) | 0.845 (3) |
| MolProbity score | 1.79 |
| Clash score | 8.11 |
| Ramachandran plot (%) |  |
| Outliers | 0.00 |
| Allowed | 2.71 |
| Favored | 97.29 |
| Rotamer outliers (%) | 1.89 |
| Cβ outliers (%) | 0.00 |
| Peptide plane (%) |  |
| Cis proline/general | 10.0/0.0 |
| Twisted proline/general | 0.0/0.0 |
| CaBLAM outliers (%) | 0.95 |
| ADP (B-factors) |  |
| Iso/Aniso (#) | 35,646/0 |
| min/max/mean |  |
| Protein | 1.58/57.10/22.25 |
| Ligand | 2.92/28.45/15.85 |
| PDB accession code | 7LMY |

**Table S3:** A summary of human p97 disease mutations and potentially affected structures

| **Disease Mutations** | **Potentially Affected Structures** |
| --- | --- |
| R93C/R93H, I114V, I151V | N domain structure |
| I126F, T127A, P137L, M158V, R159G/R159C/R159H, E185K | N-cofactor interaction |
| I27V, R95C/R95G/R95H, G97E, R155S/R155H/R155L, G156C/G156S, G157R, M158V, R159G/R159C/R159H, R191G/R191Q, L198W, G202W, A232E, T262A, K386E, N387H/N387S/N387T, A439S/A439P/A439G | N-D1 interaction |
| I206F | D1 nucleotide binding site |
| A232E | D1-D1 interaction |
| D395G | D1 domain structure |
| N401S | D1-D2 interaction |
| R487H, R662C | D2-D2 interaction |
| D592N | Pore loop II of D2 domain |
| N750S | C terminal tail |

## STAR METHODS

### Data and code availability

Cryo-EM maps have been deposited in the Electron Microscopy Data Bank (EMDB, www.ebi.ac.uk/pdbe/emdb/) under the accession codes EMDB-23449 (p97-Npl4/Ufd1-Ub_n_-Eos-CHAPSO, Class 1), EMDB-23450 (p97-Npl4/Ufd1-Ub_n_-Eos-CHAPSO, Class 2), EMDB-23451 (p97-Npl4/Ufd1-Ub_n_-Eos-CHAPSO, Class 3), EMDB-23446 (p97-Npl4/Ufd1-Ub_n_-Eos-FOM, Class 1), EMDB-23447 (p97-Npl4/Ufd1-Ub_n_-Eos-FOM, Class 2), EMDB-23448 (p97-Npl4/Ufd1-Ub_n_-Eos-FOM, Class 3), EMDB-23443 (p97-Npl4/Ufd1-Ub_6_ -FOM, Class 1), EMDB-23444 (p97-Npl4/Ufd1-Ub_6_-FOM, Class 2), EMDB-23445 (p97-Npl4/Ufd1-Ub_6_-FOM, Class 3), EMDB-23442 (p97-Npl4/Ufd1-Ub_6_-NMS-873-FOM), EMDB-23452 (p97-Npl4/Ufd1-Ub_n_-Eos-NMS-873-CHAPSO), EMDB-23453 (p97-Npl4/Ufd1-Ub_n_-Eos-NMS-873(substoichiometric)-CHAPSO, Class 1), EMDB-23454 (p97-Npl4/Ufd1-Ub_n_-Eos-NMS-873(substoichiometric)-CHAPSO, Class 2), EMDB-23455 (p97-Npl4/Ufd1-Ub_n_-Eos-NMS-873(substoichiometric)-CHAPSO, Class 3), EMDB-23456 (p97-Npl4/Ufd1-Ub_n_-Eos-NMS-873(substoichiometric)-CHAPSO, Class 4), EMDB-23457 (p97-Npl4/Ufd1-Ub_n_-Eos-NMS-873(substoichiometric)-CHAPSO, Class 5), EMDB-23458 (p97-Npl4/Ufd1-Ub_n_-Eos-NMS-873(substoichiometric)-CHAPSO, Class 6). The atomic models have been deposited in the Protein Data Bank (PDB, www.rcsb.org) under the accession codes 7LN5 (p97-Npl4/Ufd1-Ub_n_-Eos-CHAPSO, Class 1), 7LN6 (p97-Npl4/Ufd1-Ub_n_-Eos-CHAPSO, Class 2), 7LN2 (p97-Npl4/Ufd1-Ub_n_-Eos-FOM, Class 1), 7LN3 (p97-Npl4/Ufd1-Ub_n_-Eos-FOM, Class 2), 7LN4 (p97-Npl4/Ufd1-Ub_n_-Eos-FOM, Class 3), 7LMZ (p97-Npl4/Ufd1-Ub_6_ -FOM, Class 1), 7LN0 (p97-Npl4/Ufd1-Ub_6_-FOM, Class 2), 7LN1 (p97-Npl4/Ufd1-Ub_6_-FOM, Class 3), 7LMY (p97-Npl4/Ufd1-Ub_6_-NMS-873-FOM).

### Protein expression and purification

Wildtype human p97, human p97 mutants, Npl4/Ufd1, His_6_-Ub^G76V^-Ub^G76V^-mEos3.2 (Ub-Eos), human UBA1, Ube25k and gp78RING-Ube2g2 were purified as previously described^26, 32, 37^. Briefly, all proteins were expressed in *E. coli* BL21(DE3) cells and purified through Ni-NTA resin at 4 °C. All purified proteins were further buffer-exchanged to the Storage Buffer containing 50 mM HEPES, pH 8.0, 150 mM NaCl, 1 mM MgCl_2_, 0.5 mM tris(2-carboxyethyl)phosphine (TCEP) before snap freezing.

### Preparation of polyubiquitinated Ub-Eos and polyubiquitin chain

For cryo-EM studies, His-tagged Ub-Eos was directly used for polyubiquitination. For the substrate unfolding assay, Ub-Eos was first irradiated under LED UV light (385-395 nm, uvBeast V2) for 1 hour at 4 °C to induce the photoconversion of mEos3.2, followed by polyubiquitination. The polyubiquitination reaction was performed as previously described^32^. Briefly, 20 μM Ub-Eos was mixed with 1 μM UBA1, 20 μM gp78RING-Ube2g2, and 500 μM ubiquitin in the Ubiquitination Buffer containing 20 mM Hepes, pH 7.4, 150mM KCl, 10 mM ATP, and 10 mM MgCl_2_. After incubation at 37 °C overnight, the mixture was incubated with Ni-NTA resin to remove free ubiquitin chains. The elution was further separated using a Superdex 200 (GE Healthcare) size-exclusion column equilibrated in the Storage Buffer. Polyubiquitinated Ub-Eos (Ub_n_-Eos) with longer ubiquitin chains (estimated over 10 ubiquitin subunits) was collected and flash frozen for the substrate unfolding assay.

K48-linked polyubiquitin chains were assembled by mixing 1 μM UBA1, 10 μM Ube25k, and 1 mM ubiquitin in the Ubiquitination Buffer at 37 °C for 4 hours. The reaction mixture was then diluted 10 folds with 50 mM NaOAc, pH 4.5, and separated by a Mono S cation exchange column. The corresponding peak for hexa-ubiquitin chain (Ub_6_) was collected and further purified with a Superdex 75 (GE Healthcare) size-exclusion column equilibrated in the Storage Buffer.

### Assembly of substrate-engaged p97 complex

p97-Npl4/Ufd1 complex was assembled as previously described^26^. A p97 mutant bearing A232E and E578Q mutations was used in order to decrease the heterogeneity of N domains and reduce the unfolding activity of the complex. Two-fold molar excess of Ub_6_ or Ub_n_-Eos was added to the p97-Npl4/Ufd1 complex, followed by gel filtration using a Superose 6 column (GE Healthcare) equilibrated in the Storage Buffer (**Extended Data Fig. 1b and 1c**). The assembled substrate-engaged p97 complex was flash frozen in liquid nitrogen for structural studies. No nucleotide was supplemented during the assembly.

### Specimen preparation for single-particle cryo-EM

All samples for cryo-EM were prepared as previously described^26^, with the following modifications. Either fluorinated octyl maltoside (FOM) or CHAPSO was used to relieve the preferred orientations instead of IGEPAL CA-630. The complex was concentrated to ∼2.5 mg/mL followed by incubation with 5 mM ATP at room temperature for 5 minutes before adding FOM (final concentration 0.01% v/v). Glow discharged Quantifoil Cu 1.2/1.3 grids were used for FOM supplemented samples. Alternatively, the samples were concentrated to ∼20mg/mL followed by incubation with 5 mM ATP at room temperature for 5 minutes before adding CHAPSO (final concentration 4 mM). Non-glow-discharged Quantifoil Cu 1.2/1.3 grids were used for CHAPSO supplemented samples. In both cases, 3.5 μL sample was applied to the grid and was blotted for 1 second using standard Vitrobot filter paper (Ted Pella, 47000-100) before plunge freezing in liquid ethane. For substrate-engaged p97-Npl4/Ufd1 complex in the presence of NMS-873, besides the aforementioned procedures, NMS-873 was supplemented right after the incubation with ATP at either 10 μM or 80 μM final concentration. The samples were incubated for another 30 min at room temperature before adding the detergents and plunge freezing.

### Data collection for single-particle cryo-EM

Data collection was performed either in Advanced Electron Microscopy Facility at the University of Chicago or National Cryo-Electron Microscopy Facility at the National Cancer Institute. All datasets were acquired as movie stacks with a Titan Krios electron microscope operating at 300 kV, equipped with either a Gatan K2 Summit or K3 direct detector camera. A single stack typically consists of 40 frames with a total exposure around 50 electrons/Å**^2^**. The defocus range was set at −1.0 to −2.5 μm. See **Supplementary Table 1** for the details.

### Image processing

Movie stacks were subjected to beam-induced motion correction using MotionCor2^38^. CTF parameters for each micrograph were determined using CTFFIND4^39^. Particle picking, two- and three-dimensional classifications, three-dimensional refinement, and local resolution estimation were performed in RELION-3^40^. Particle picking was performed by manually choosing ∼2,000 particles and generating templates through reference free 2D classification, followed by automatic template based picking. False-positive particles or particles classified in poorly defined classes were discarded after 2D classification. The initial 3D classification was performed on a binned dataset with the previously reported p97 structures as the reference model^26^. The detailed data processing flows are shown in Figures S2, 3, 4, 9, 10, and 11. To make sure that the 3D classification did not miss any major conformations, additional runs were performed with different number of classes (ranges from 3-8) and different regularization parameters (ranges from 2-6). Since we did not observe a dependence on the regularization parameter, the results from the default value (T=4) were shown in the figures. Data processing statistics are summarized in **Supplementary Table 1**. Reported resolutions are based on Fourier shell correlation (FSC) using the FSC=0.143 criterion.

### Model building, refinement, and validation

Model building was based on the existing cryo-EM structures of human p97^26^ (PDB ID: 7JY5). The models of individual domains (N, D1, and D2) were first docked into the cryo-EM maps as rigid bodies using UCSF Chimera^41^ followed by further adjustment using COOT^42^. For well-resolved regions, sharpened maps were used and individual residues were manually adjusted and fit into the density. For flexible regions, unsharpened maps were used and individual secondary structure elements or subdomains were fit into the density as rigid bodies. A short β strand was automatically built into the density corresponding to the translocating peptide using COOT, followed by manual extension on either ends and real-space refinement until the all the density was covered. The final models were subjected to global refinement and minimization in real space using the real-space refinement module in Phenix^43^. Model validation was performed using the comprehensive validation tool in Phenix. The statistics of model refinement is shown in **Supplementary Table 2**.

### Substrate unfolding assay

Substrate unfolding assay was performed as previously described^26, 32^. Briefly, 20 nM photoconverted polyubiquitinated Ub-Eos was mixed with 400 nM p97 or p97 mutants and 500 nM Npl4/Ufd1 in the Assay Buffer containing 50 mM Tris pH 7.4, 5 mM KCl, 20 mM MgCl_2_, 1 mM EDTA, 0.5 mM TCEP, and 0.01% Triton. Proteins were pre-incubated in a 96-well plate (Costar 3694) for 10 minutes at 37 °C before initiating the reaction by supplementing the ATP regeneration mixture (5 mM ATP, 30 mM creatine phosphate, and 50 μg/mL creatine phosphokinase). Fluorescence signal was monitored using a TECAN safire2 plate reader at 540 nm excitation and 580 nm emission wavelengths with 30 seconds intervals for 60 min. Each reaction condition was repeated three times. Background fluorescence was measured by mixing the same amount of substrate with 6 M guanidine-HCl and was subtracted from the average of the experimental groups. Normalized fluorescence was plotted using GraphPad Prism 8.4.2.

